# Attentive graph neural network models for the prediction of blood-brain barrier permeability

**DOI:** 10.1101/2024.10.12.617907

**Authors:** Jesse W. Collins, Mahmoud Ebrahimkhani, Daniel Ramirez, Jonathan Deiloff, Mauro Gonzalez, Mostafa Abedi, Laurence Philippe-Venec, Bridget M. Cole, Brandon Moore, Jennifer O. Nwankwo

## Abstract

The blood-brain barrier’s (BBB) unique endothelial cells and tight junctions selectively regulate passage of molecules to the central nervous system (CNS) to prevent pathogen entry and maintain neural homeostasis. Various neurological conditions and neurodegenerative diseases benefit from small molecules capable of BBB penetration (BBBP) to elicit a therapeutic effect. Predicting BBBP often involves *in silico* assessment of molecular properties such as lipophilicity (log*P*) and polar surface area (PSA) using the CNS multiparameter optimization (MPO) method. This study curated an open-source dataset to rigorously benchmark machine learning (ML) and neural network (NN) models with each other and with MPO for predicting BBBP. Our analysis demonstrated that AI models, especially attentive NNs using stereochemical features, significantly outperform MPO in predicting BBBP. An attentive graph neural network (GNN), which we refer to as CANDID-CNS™, achieved a 0.23-0.26 higher AUROC score than MPO on full test sets, and a 0.17-0.19 higher score on stereoisomers filtered subsets. Regarding stereoisomers that differ in BBBP, which MPO cannot distinguish, attentive GNNs correctly classify these with AUROC and MCC metrics comparable to or better than MPO’s AUROC and MCC on less difficult test molecules. These findings suggest that integrating attentive GNN models into pharmaceutical drug discovery processes can substantially improve prediction rates, and thereby reduce the timeline, cost, and increase the probability of success of designing brain penetrant therapeutics for the treatment of a wide variety of neurological and neurodegenerative diseases.

## 1. Introduction

The blood-brain barrier (BBB) is a highly selective, permeable anatomical structure critical for maintaining neural homeostasis and protecting the CNS from harmful substances^1,2^. Here, we use the terms BBB and CNS permeability interchangeably, noting that the most likely access route of small molecules to the CNS is through the large surface area of the BBB^3^. Anatomically, the BBB consists of brain endothelial cells lining the cerebral microvessels, which are distinct from peripheral endothelial cells due to their unique tight junctions^2,4^. These junctions create a physical barrier that restricts paracellular transport, thereby controlling the ionic balance of the brain and shielding it from toxins and pathogens^5^. The endothelial cells exhibit minimal pinocytic activity and have specialized transport systems, including efflux transporters like P-glycoprotein (P-gp)^6^. These systems actively pump certain molecules out of the brain and selectively allow essential nutrients, such as glucose and amino acids, to enter^7^.

Certain pathogens can compromise the integrity of the BBB, facilitating the entry of bacteria or viruses and leading to conditions such as meningitis and encephalitis^8,9^. Neuroinflammatory diseases such as multiple sclerosis (MS), and neurodegenerative disorders such as Alzheimer’s Disease (AD) and Parkinson’s Disease (PD) often involve a compromised BBB. In MS, BBB breakdown permits immune cells to invade the brain, attacking the myelin sheath and leading to inflammation and neuronal damage^10^. This process necessitates developing immunomodulatory drugs that can cross the BBB to modulate or suppress the abnormal immune activity within the CNS^11,12^. Similarly, in AD and PD, diseases which are characterized by accumulation of amyloid beta plaques and dopaminergic neurodegeneration, respectively, therapeutic efficacy depends on the drug’s ability to cross the BBB^13,14^. Additionally, effective management of brain tumors, including glioblastoma, requires chemotherapeutic agents to penetrate the BBB^15^.

This intricate anatomy and physiology of the BBB makes the delivery of drugs to the CNS challenging. Small molecule therapeutics present a promising approach for alleviating symptoms and slowing CNS disease progression. Predicting the effectiveness of such agents requires a detailed understanding of chemical and physiological properties such as lipophilicity and polar surface area (PSA) that influence BBB permeability (BBBP)^16–18^. Balancing these properties is essential not only for effective BBBP but also for ensuring adequate solubility, absorption, and distribution. Moreover, the interaction of a molecule with transport proteins and efflux systems at the BBB, such as P-gp, is paramount^19^. Recently, a proteomics informed relative expression factor (REF) method in conjunction with brain PET imaging has demonstrated effective prediction of the brain-to-plasma concentration ratio of drugs expelled by P-gp efflux^20^.

Pharmacokinetic (PK) studies that determine CNS exposure precede *in vivo* efficacy and clinical studies but can be costly and time consuming. Consequently, researchers typically prioritize compounds for *in vivo* studies based on their *in silico* predicted likelihood of achieving adequate CNS exposure. However, this reliance on *in silico* models poses a risk of overlooking BBBP positive (BBB+) compounds due to potential model inaccuracies, a significant concern considering that only about 2% of small molecule therapeutic candidates cross the BBB^21^.

BBBP prediction is broad phrasing that warrants further contextualization. PK studies that determine whether a compound crosses the BBB typically involve administration of the compound to mice and subsequent measurement of the concentration of the compound in the brain and the blood at multiple time points. These measurements provide K_p,brain_, the brain to plasma partition coefficient, reflecting the ratio of the total concentration of the compound in the brain to its total concentration in the blood^3^. A major improvement upon this measurement is K_p,uu_^22^, the unbound brain to unbound blood plasma partition coefficient, which can be estimated by measuring K_p,brain_ and the unbound drug fractions via *in vitro* brain tissue and plasma protein binding assays^3,22^. These three measurements can be used together to estimate the ratio of freely available drug in the brain to that in the bloodstream, a more pharmacologically relevant parameter than the total concentration ratio^3,22^.

Although quantitative measurements of BBBP are vital for CNS drug discovery programs, the focus of our study is on classification of potential CNS candidates rather than prediction of numerical measurements of BBBP such as K ^23,24^. The reasons for this modeling focus are threefold. 1) A state-of-the-art CNS drug discovery method, multiparameter optimization (MPO),^25^ already exists for classification purposes, and serves as the primary benchmark for the ML and NN approaches we study here. 2) Compared to a regressor’s numerical output, the 0 to 1 probability output of a classification model can be applied more straightforwardly in reinforcement learning for guiding molecule generation AIs toward producing potential hits with favorable absorption, distribution, metabolism, excretion, and toxicity (ADMET) properties, such as BBBP in the case of CNS therapeutics. 3) The most thoroughly curated public data relevant to this ADMET endpoint is primarily based on site of action clinical and post-clinic findings^26^. Primary curation studies typically infer that drugs with CNS activity are BBB+, while drugs with a non-CNS site of action and without known CNS side effects may be categorized as BBB-by expert medicinal chemists and pharmacologists^27–29^. This curation process enables the use of many public compounds for modeling CNS and non-CNS drugs without quantitative measurements of BBBP. The high opportunity cost of missing a BBB+ compound is underscored by instances where neuroscience therapeutics candidates failed to demonstrate sufficient brain permeability in clinical evaluations. For example, Alzheimer’s disease drug candidate AB-957 (Alicapistat) lacked clinical efficacy on REM sleep patterns^30^. This is most likely attributed to low CNS exposure since drug concentrations in cerebrospinal fluid (CSF) of clinical trial subjects were below its *in vitro* calpain inhibition constant (Ki)^30^.

In addition, it is well established that the enantiomers of the pain drug ibuprofen show variations in CSF concentrations, potentially due to stereoselective metabolic processes^31,32^. Not only metabolism, but also active transport^33–35^ and P-gp efflux^36–38^ can affect free fraction across the BBB in a stereoselective manner. Moreover, the tragedy of thalidomide^39^ demonstrates the need to consider stereochemistry in all small molecule drug discovery programs. Ideally, *in silico* BBBP prediction models would be sensitive to stereoisomerism to accurately assess CNS penetration potential.

Recent advances in *in silico* BBBP prediction have included the development of larger public datasets^26^ and the application of conventional machine learning (ML)^40^, neural networks (NNs)^40–43^, and physics^44^ methods. These modern approaches contrast with the established MPO method^25^, which assesses BBBP and other drug characteristics through molecular weight (MW), hydrogen bond donors (HBD), topological polar surface area (TPSA), calculated lipophilicity (clog*P*), and calculated distribution coefficient (clog*D*)^45,46^. This study examines the inclusion and weighting of these parameters in MPO, especially in comparison to pretrained NN models finetuned for BBBP prediction.

Prior studies have indicated that ML and NN models, even those trained on datasets of approximately 2,000 small molecules, may surpass MPO for predicting BBBP^47,48^. Building on these findings, our study employs a larger, more meticulously curated dataset to explore the unique advantages of BBBP classifiers. We particularly focus on the capabilities of conventional ML and NN models to classify compounds such as those with intramolecular hydrogen bonds (IMHBs)^49^, or stereoisomers of differing BBBP, which MPO might not identify as BBB+ compounds correctly. Our analysis extends to the highly curated B3DB dataset^26^, which we further refined to include drug-like molecules and a common set of molecules amenable to cross-validation across various AI/ML architectures.

These AI/ML architectures, chosen for their diversity in input features, range from calculated descriptors to images, graphs, and multiple 3D conformations, include Attentive FP GNN (AttFP GNN)^50^, support vector classifier (SVC)^51^, XGBoost classifier (XGBC)^52^, logistic regressor (LR), and pretrained models like MolCLR^53^, ImageMol^54^, and Uni-Mol^55^. By applying a more rigorous and statistically meaningful cross-validation process^56^, with particular attention to stereoisomers of varying BBBP class, our study reveals which features, pretraining strategies, and model architectures are most suitable for classifying brain penetrant compounds. By benchmarking the performance of these models on an internal BBBP dataset and assessing Tanimoto similarities, our study also aims to understand the extent to which molecular similarity to training data influences model performance. Finally, we discuss how our results could be used to improve MPO, as well as further considerations regarding potency and toxicity when evaluating the usage of an ML or NN BBBP model over MPO.

Our results suggest that AttFP GNN models, such as CANDID-CNS*™*, highlighted here, can supplant traditional MPO methodologies for BBBP prediction. Integrating attentive GNN models into pharmaceutical drug discovery processes can significantly improve prediction rates, reduce costs and timelines, and increase the probability of discovering brain penetrant therapeutics for the treatment of a wide variety of neurological and neurodegenerative diseases.

## 2. Methods

### 2.1. B3DB data curation and analysis

Our study involved a meticulous curation process of the B3DB dataset to ensure data quality and relevance for different ML and NN models. Initially, we filtered out compounds with missing or invalid SMILES (Simplified Molecular Input Line Entry System) strings and those representing molecules with unusual charge characteristics. Further refinement involved preprocessing the molecular structures, including selecting the freebase form, standardizing structures, and removing duplicates, with the final dataset comprising 7465 molecules. Detailed steps of this curation process, including specific criteria for removal and preprocessing techniques, are provided in Supplementary Information.

For our MPO analysis that follows curation, we compared MPO parameters on the filtered dataset by their BBB± category, and also use 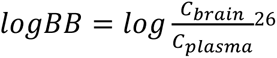, for a quantitative analysis, where ^*C*^*_brain_* is total concentration of the molecule in the brain and *C_plasma_* is the total concentration of the molecule in the blood. This measurement is not as rigorous as K_p,uu_, but we hypothesize that the trends we observe in log*BB* would likely hold for K_p,uu_ measurements as a function of these MPO parameters as well.

### 2.2. Dataset splitting for cross validation

We employed scikit-learn^51^ and in house scripts to conduct 5 fold cross-validation on the dataset. The data was divided into training and test sets with an 80:20 ratio. For models utilizing a validation set for early stopping, the test set was further split 50:50, resulting in an overall distribution of 80:10:10 across training, validation, and test sets, respectively. In contrast, for models like SVC and XGBC that do not require a validation set, we adopted a 90:10 training to test set split. Our cross-validation approach included both random and stratified splits. We also report results with a single scaffold split that factored in chirality during scaffold assignment.

### 2.3. Stereoisomer Filtering

For a more rigorous evaluation of BBBP models, we prepared subsets of our test sets, specifically focusing on stereoisomer content. Our processed dataset of 7465 molecules was grouped by nonisomeric SMILES to identify stereoisomer groups, categorizing them as either entirely BBB+, entirely BBB-, or mixed (“multiclass”).

For more stringent testing, we removed stereoisomers from the test set if they belonged to a single class group and had a counterpart in the corresponding training or validation set for that fold. However, stereoisomers in multiclass groups were retained in the test set, regardless of their presence in training or validation sets, to assess the classifiers’ ability to generalize and predict BBBP class based on structural features rather than training memorization.

The performance comparison was then focused exclusively on these multiclass stereoisomer subsets, providing insights into the models’ capacity to differentiate stereoisomers with varying BBBP classes. Additionally, a separate analysis was conducted to evaluate the role of stereoisomers in BBBP model development, with details of this procedure and its outcomes available in the Supplementary Information.

### 2.4. Feature extraction

We employed the Mordred molecular descriptor calculator^57^ to calculate approximately 1000 descriptors for each molecule. This was part of our initial phase in training a preliminary XGBoost model, following the train and test sets defined by Roy et al.^44^, with the primary goal of determining feature importance. Based on this analysis, we selected the top 50 Mordred descriptors, with the TPSA emerging as the most significant. These selected descriptors were then utilized in the cross-validation training of XGBoost and SVC models.

For the AttFP GNN model, CANDID-CNS*™*, we generated molecular graphs using PyTorch Geometric^58^, incorporating one-hot encoding for chiral atoms and bonds to enhance the model’s ability to recognize molecular chirality. Regarding the models ImageMol, MolCLR, and Uni-Mol, we processed features using their respective codebases. Specifically for Uni-Mol, we aimed to generate 10 3D conformations for each molecule within our dataset. However, 14 molecules failed to embed in 3D in each of the 10 attempts. In such instances, we used 2D coordinates instead, ensuring continuity in our analysis.

### 2.5. Model training and hyperparameter tuning

We employed various model architectures, each with specific hyperparameters and training mechanisms to optimize their performance. For the SVC^51^, we used scikit-learn’s implementation with a regularization parameter of 100 and a gaussian radial basis function kernel. For XGBoost, we determined the best hyperparameters through a grid search, including a learning rate of 0.05 and max depth of 20. The AttFP GNN was adapted into CANDID-CNS™ as a classifier with a sigmoid layer, optimized using the Adam optimizer with a learning rate of 10^-2.5^. We also explored a chiENN version of AttFP GNN with adjusted hyperparameters to handle chiENN edge graphs.

ImageMol, MolCLR (graph convolution network (GCN) and graph isomeric network (GIN), and Uni-Mol models were finetuned from their pretrained models with default or modified settings for epochs, learning rate, and dropout ratio. The detailed hyperparameters for each model and the specific training procedures are provided in the Supplementary Information. All models benefiting from GPU acceleration were trained using NVIDIA V100s on Microsoft Azure ML Studio.

### 2.6. Performance evaluation

In evaluating our classifiers, we utilized standard metrics including sensitivity, area under the receiver operating characteristic curve (AUROC), accuracy, and Matthews correlation coefficient (MCC). Sensitivity was emphasized due to the significant opportunity cost associated with failing to identify a BBB+ compound in neuroscience drug discovery. Additionally, we reported the AUROC, accuracy, and MCC, as they provide a more comprehensive assessment of the classifier’s overall performance. Precision and specificity were also considered to ensure that a high sensitivity value did not result from an excessive number of false positives. The exact equations and definitions for each of these parameters are detailed in the Supplementary Information.

### 2.7. *In vivo* brain permeability measurements

For internal compound BBBP data, we studied an *in vivo* mouse model of brain and plasma compound exposure determined in male and female mice following intravenous administration of compound at a dose of 3 mg/kg, with data for each gender determined in triplicate. We used log*BB*, with a threshold of −1, to distinguish internal compound classes due to the absence of *in vivo* efficacy and plasma protein binding for most compounds in this set.

### 2.8. MPO calculation

We adopted the approach described by Wager et al.^46^ for scoring various molecular properties. This involved using monotonic decreasing transformations for MW, clog*P*, clog*D*, most basic pKa, and HBD values and a hump transformation for scoring the TPSA values. Monotonic decreasing transformations assign higher scores to lower values of a parameter, decreasing towards zero for higher values, while the hump transformation assigns the highest scores to an intermediate range of values, tapering off to zero for both lower and higher extremes.

For the calculation of TPSA, we utilized the RDKit implementation^59^, based on the work by Ertl et al^60^. This calculation was modified to include sulfur and phosphorus atoms, acknowledging their prevalence in our dataset and their observed influence on the MPO score. The cLogP values were also derived using the RDKit implementation, following the methodology of Wildman and Crippen validated against nearly 10000 measurements determined by Biobyte^61^. The most basic pKa value was calculated using MolGpKa^62^, a GNN based pKa calculator. For clog*D* values within our MPO model, we referred to our internal model, which is described in the next section. Additionally, we separately reported MPO scores generated by the CDD Vault software, provided by Collaborative Drug Design (Burlingame, California), allowing for an alternative perspective on the MPO scoring of compounds.

### 2.9. Lipophilicity (logD) models and measurements

In assessing lipophilicity, we utilized two distinct methods. The standard shake flask method quantifies compound concentration in two immiscible solvents, such as octanol/water, suitable for a lipophilicity range of −1 to 2.8. For lipophilicity measurements beyond this range, we employed the Chromlog*D* method, which operates on a lipophilic C18 support at different pH to determine the Chromatographic Hydrophobicity Index (CHI). CHI provides insights into the compound’s distribution between stationary and mobile phases and its ionization stage. Our high performance liquid chromatography (HPLC) protocol for Chromlog*D*, in collaboration with Aurigene Services Limited (Hyderabad, Andra Pradesh, India), was rigorously standardized using chromatographic standards^63^.

We developed two models to calculate SFlog*D* and Chromlog*D*. Our AttFP GNN model for calculated clogD was trained on a large dataset of 1.4 million ChemAxon clog*D* values^64^, supplemented with proprietary data on internal compounds. Training of our calculated Chromlog*D* model, a variant of the AttFP model^65^, included a dataset of clog*D* ChemAxon values rescaled to Chromlog*D* scale using dual log*D*/Chromlog*D* measurements. Both models were optimized for performance using specific hyperparameters, learning rates, and training epochs. Detailed specifications of these models are available in the Supplementary Information.

## 3. Results and Discussion

### 3.1. MPO Analysis

Here, we investigate the impact of various parameters within MPO for BBBP prediction. We start with a categorical analysis, then focus on the correlation of each MPO parameter with measured log *BB* values. A subset of 991 compounds, or about 15% of the total using for training, validation, and testing, had associated log*BB* values within the dataset^26^. Although log*BB* is related to total brain to plasma concentration, and not as meaningful as K_p,uu_, we work with this data for two reasons beyond it being more abundant than K_p,uu_ data: 1) We find that log*BB* values tend to correlate strongly with K_p,uu_ values that are shared in the literature, and 2) we found that our models, which make use of the 85% of compounds that are classified due to site of action information alone, are also predictive of K_p,uu_ (**Figure S3**).

Individual MPO components do not distinctly differentiate between BBB+ and BBB-classes (**Figure S4**). However, certain thresholds, such as a TPSA of less than 120, fewer than 3 HBD, MW below 500, and values of clog*P* and clog*D* greater than 1 (**Figure S4a**), demonstrate some potential in identifying BBB+ compounds while minimizing false negatives. Notably, TPSA exhibits the most significant Spearman correlation or anticorrelation (−0.59) with measured log*BB* values (**Figure S4b)**, highlighting its pivotal role in MPO for BBBP prediction, despite being weighted equally with other components in the MPO score. Both clog*P* and clog*D* correlate with log*BB*, suggesting that their respective MPO scoring functions, which penalize high clog*P* or clog*D* due to toxicity concerns, may inadvertently limit MPO’s effectiveness for BBBP prediction. Additionally, we observed that most MPO component values tend to exhibit correlation or anticorrelation with one another (**Figure S5**). Given their lack of linear independence, the approach of scoring and summing these components in MPO could result in limited enhancement for BBBP classification.

To further explore the impact of MPO parameters BBBP predictions, we trained a logistic regressor (LR) using the individual component values of MPO, rather than their aggregated scores, to predict the BBBP class. Notably, TPSA emerged as the most influential factor in the LR model for predicting BBBP, as depicted in **(Figure S6a)**. This finding aligns with report on an XGBoost model trained with another BBBP dataset^41^, where TPSA’s importance is similarly highlighted. In our XGBoost model’s use of Mordred descriptors, TPSA also holds the most significance in BBBP prediction. Following TPSA, the LR model showed that clog*P* had a slight influence on probability outcomes, with other MPO components contributing minimally **(Figure S6a)**. The coefficients of the LR, as shown in **(Figure S6b)**, predominantly correlate or anticorrelate with TPSA. This pattern suggests that the equal weighting of these components in the traditional MPO scoring system may not effectively improve its utility for BBBP classification. Given the prominence of TPSA and clog*P* in the LR model, we further explored their standalone use as classifiers. Specifically, we assessed the performance of classifiers based solely on TPSA, and on a combination of TPSA and clog*P* scores, to determine their utility in BBBP class prediction. The results of this analysis are reported in **Figures 1-2** and **Tables 1-2**.

**Figure 1.**
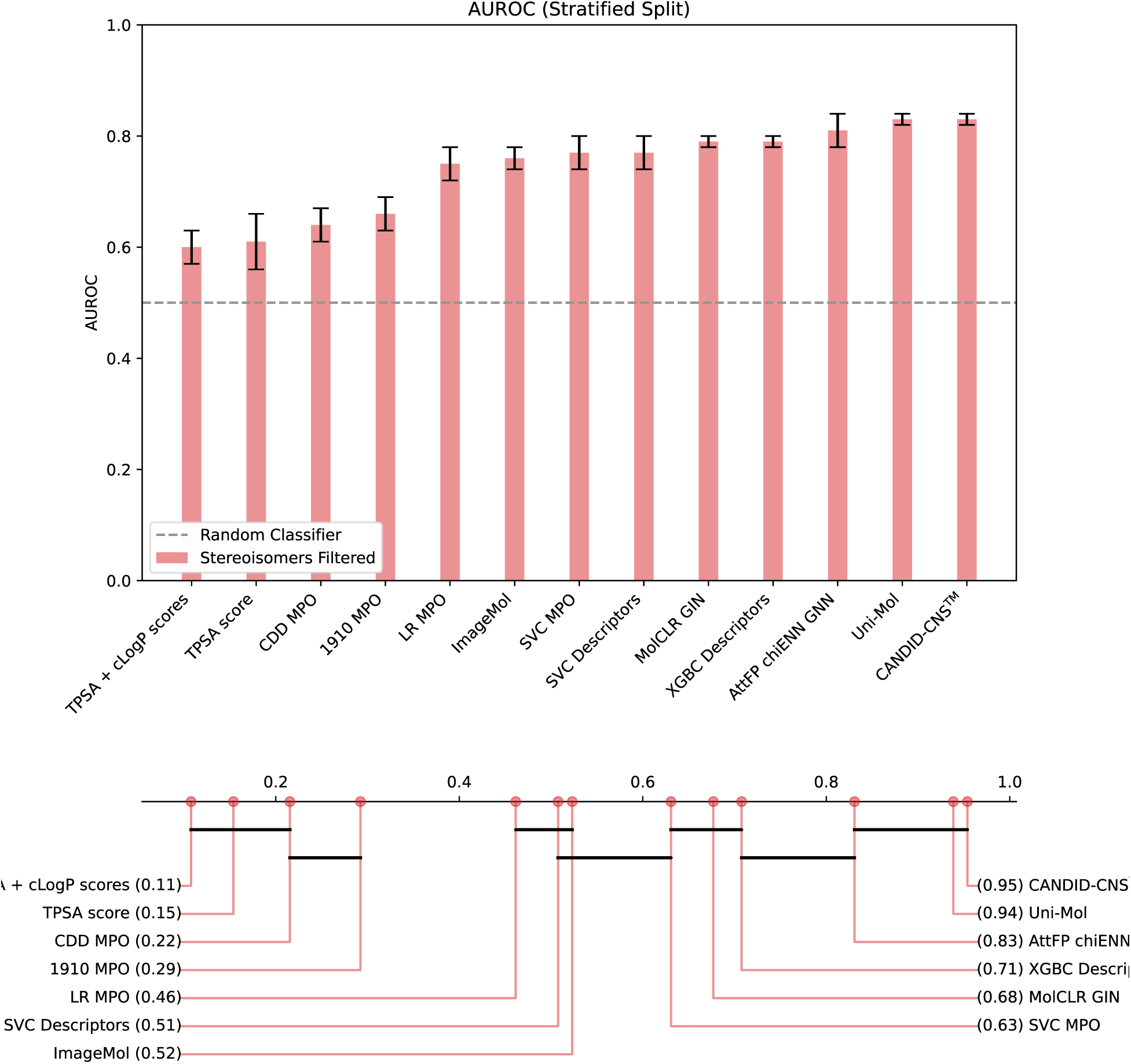
CANDID-CNS™ Leads in AUROC All Models When Removing Single Class Stereoisomers From Test Sets. Error bars represent 1 standard deviation from mean. a) AUROC results show CANDID-CNS™ performs best. b) Critical difference diagram showing CANDID-CNS™, Uni-Mol, and the Attentive FP GNN with chiENN layer all perform statistically better than MPO by Friedman rank. Models are connected if pairwise Conover p-value is > 0.05.

**Figure 2.**
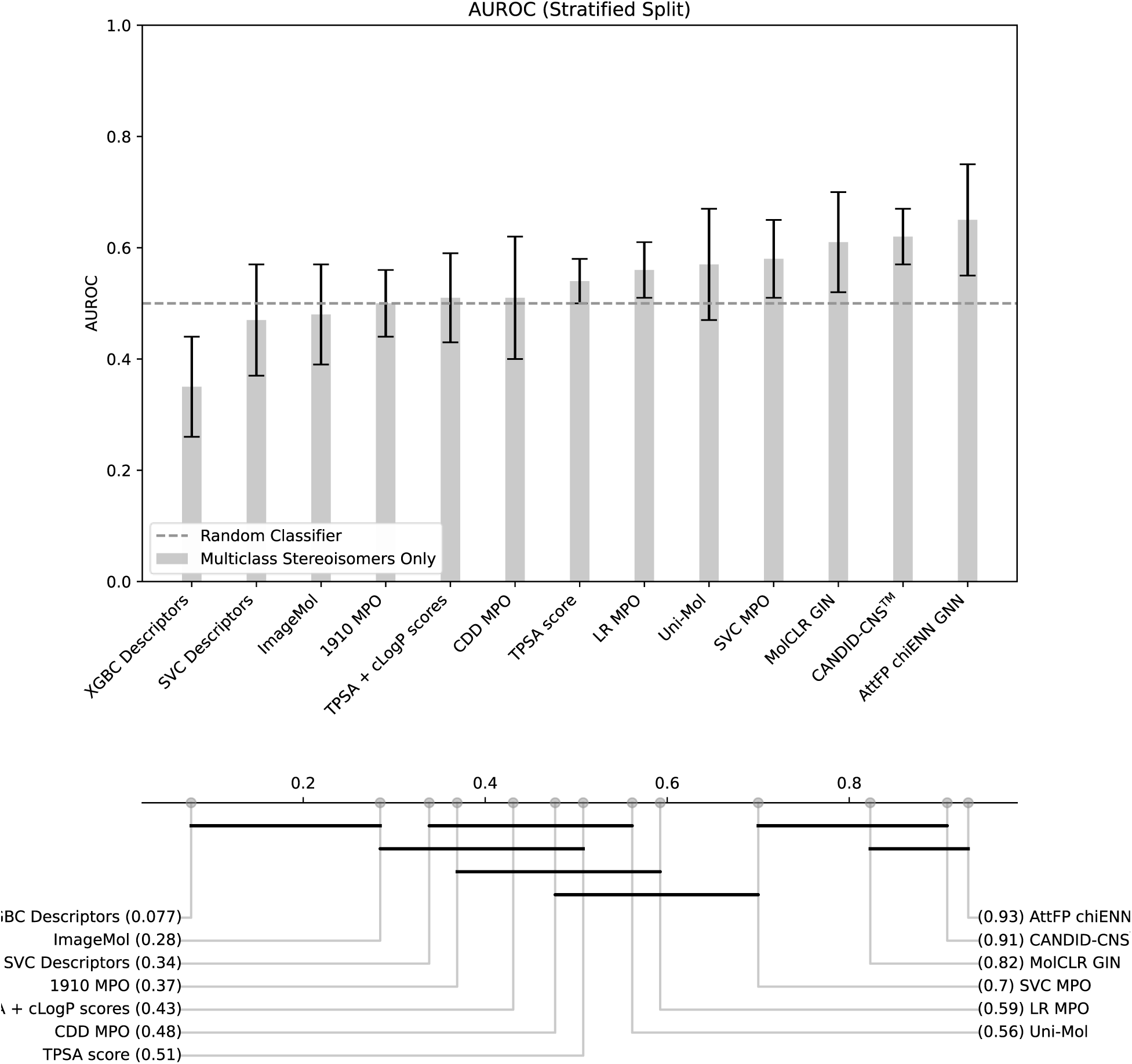
CANDID-CNS™ and chiENN Version Perform Best in Distinguishing BBBP Class Among Stereoisomers. a) AUROC results on multiclass stereoisomers in the test sets show CANDID-CNS™ and the chiENN version performing best. Error bars represent 1 standard deviation from mean. b) Critical difference diagram suggesting these models significantly outperform MPO and other models in AUROC on this task. Models are connected if pairwise Conover p-value is > 0.05.

**Table 1.** Model Comparison on Stereoisomers Filtered Test Subsets.

| Classifier (Stratified) | AUROC | MCC | AUPRC | Sensitivity | Accuracy | Precision | Specificity |
| --- | --- | --- | --- | --- | --- | --- | --- |
| Uni-Mol | **0.83±0.01** | **0.53±0.03** | **0.88±0.02** | **0.78±0.09** | **0.77±0.02** | 0.83±0.04 | 0.76±0.09 |
| CANDID-CNS™ | **0.83±0.01** | **0.53±0.01** | 0.87±0.02 | 0.73±0.05 | 0.76±0.01 | 0.85±0.03 | 0.81±0.05 |
| AttFP chiENN GNN | 0.81±0.03 | 0.50±0.05 | 0.86±0.02 | 0.73±0.09 | 0.75±0.03 | 0.83±0.03 | 0.77±0.08 |
| MolCLR GIN | 0.79±0.01 | 0.48±0.02 | 0.84±0.03 | 0.64±0.07 | 0.72±0.02 | **0.86±0.04** | **0.84±0.07** |
| XGBC Descriptors | 0.79±0.01 | 0.48±0.03 | 0.85±0.02 | 0.70±0.10 | 0.73±0.03 | 0.83±0.04 | 0.80±0.04 |
| SVC MPO | 0.77±0.03 | 0.45±0.05 | 0.81±0.01 | 0.69±0.08 | 0.72±0.03 | 0.81±0.02 | 0.76±0.05 |
| SVC Descriptors | 0.77±0.03 | 0.43±0.05 | 0.82±0.03 | 0.68±0.06 | 0.71±0.03 | 0.81±0.03 | 0.76±0.05 |
| ImageMol | 0.76±0.02 | 0.43±0.05 | 0.81±0.01 | 0.72±0.06 | 0.72±0.02 | 0.79±0.03 | 0.72±0.08 |
| LR MPO | 0.75±0.03 | 0.41±0.06 | 0.79±0.01 | 0.71±0.05 | 0.71±0.03 | 0.78±0.02 | 0.78±0.08 |
| 1910 MPO | 0.66±0.03 | 0.29±0.04 | 0.72±0.03 | 0.75±0.02 | 0.67±0.02 | 0.71±0.02 | 0.54±0.05 |
| CDD MPO | 0.64±0.03 | 0.27±0.04 | 0.71±0.02 | 0.64±0.17 | 0.63±0.05 | 0.73±0.03 | 0.63±0.14 |
| TPSA score | 0.61±0.05 | 0.22±0.08 | 0.66±0.03 | 0.75±0.09 | 0.63±0.03 | 0.67±0.03 | 0.46±0.10 |
| TPSA + cLogP scores | 0.60±0.03 | 0.23±0.07 | 0.67±0.02 | 0.61±0.16 | 0.61±0.05 | 0.71±0.02 | 0.62±0.11 |

**Table 2.** Model Performance on Multiclass Stereoisomer Test Subsets. Data splitting is stratified unless otherwise specified.

| Classifier | AUROC | MCC | AUPRC | Sensitivity | Accuracy | Precision | Specificity |
| --- | --- | --- | --- | --- | --- | --- | --- |
| AttFP chiENN GNN | **0.65±0.10** | **0.38±0.12** | 0.61±0.11 | 0.80±0.16 | 0.66±0.07 | 0.61±0.14 | 0.58±0.24 |
| CANDID-CNS™ | 0.62±0.05 | 0.34±0.09 | 0.58±0.09 | 0.59±0.26 | 0.63±0.05 | 0.65±0.19 | 0.72±0.20 |
| MolCLR GIN | 0.61±0.09 | 0.37±0.15 | 0.60±0.08 | 0.71±0.32 | 0.69±0.06 | 0.64±0.08 | 0.60±0.28 |
| SVC MPO | 0.58±0.07 | 0.25±0.08 | 0.52±0.08 | 0.63±0.20 | 0.62±0.06 | 0.56±0.09 | 0.61±0.16 |
| Uni-Mol | 0.57±0.10 | 0.29±0.14 | 0.54±0.08 | 0.70±0.29 | 0.64±0.10 | 0.66±0.20 | 0.53±0.32 |
| LR MPO | 0.56±0.05 | 0.24±0.08 | 0.51±0.10 | 0.68±0.13 | 0.62±0.05 | 0.54±0.09 | 0.56±0.07 |
| TPSA score | 0.54±0.04 | 0.16±0.08 | 0.46±0.09 | 0.58±0.09 | 0.58±0.05 | 0.52±0.08 | 0.58±0.12 |
| CDD MPO | 0.51±0.11 | 0.23±0.14 | 0.50±0.03 | 0.36±0.28 | 0.61±0.12 | N/A | 0.86±0.12 |
| TPSA + cLogP scores | 0.51±0.08 | 0.18±0.13 | 0.48±0.06 | 0.37±0.19 | 0.61±0.09 | N/A | 0.80±0.15 |
| 1910 MPO | 0.50±0.06 | 0.24±0.09 | 0.51±0.06 | 0.45±0.21 | 0.63±0.07 | 0.69±0.18 | 0.75±0.21 |
| ImageMol (Scaffold) | 0.50 | 0.18 | **0.72** | 0.53 | 0.57 | **0.80** | 0.67 |
| ImageMol (Random) | 0.49±0.05 | 0.22±0.06 | 0.46±0.05 | 0.73±0.31 | 0.58±0.04 | 0.53±0.06 | 0.44±0.27 |
| ImageMol | 0.48±0.09 | 0.18±0.11 | 0.47±0.14 | 0.62±0.28 | 0.54±0.07 | 0.57±0.23 | 0.53±0.34 |
| SVC Descriptors | 0.47±0.10 | 0.23±0.15 | 0.45±0.09 | 0.52±0.39 | 0.61±0.08 | 0.62±0.22 | 0.67±0.26 |
| Uni-Mol (Random) | 0.44±0.07 | 0.19±0.11 | 0.42±0.08 | 0.74±0.38 | 0.56±0.05 | N/A | 0.39±0.32 |
| Uni-Mol (Scaffold) | 0.43 | 0.35 | 0.70±0.00 | **1.00** | **0.76** | 0.75 | 0.17 |
| XGBC Descriptors | 0.35±0.09 | 0.06±0.09 | 0.38±0.07 | 0.14±0.26 | 0.56±0.10 | N/A | **0.91±0.16** |

Our analysis of the distribution of MPO components and Chromlog*D* values for internal compounds **(Figure S7)** indicates that the current literature definition of MPO is suboptimal for predicting BBBP in these molecules. We observed that our BBB+ compounds exhibit higher average clog*D* and Chromlog*D* values than BBB- compounds, which contrasts with the typical weighting of clog*D* in MPO. While MPO scores, particularly for clog*P* and clog*D*, may act as safeguards against highly lipophilic compounds with potential off target effects, they appear less effective for BBBP prediction. In the LR model trained on processed B3DB data, clog*P* demonstrates a diminishing weight and fails to adequately distinguish between our BBB+ and BBB-compounds (**Figure S6**). HBD significantly differentiates between these classes, whereas MW does not show such a distinction **(Figure S7)**. Predicted values of the most basic pKa also show some differentiation between the classes **(Figure S7)**, but all predicted pKa values would be weighted similarly in the MPO parameter scoring functions from the published 2016 study we utilized^48^.

### 3.2. Stereoisomer Analysis

In our analysis of the 7465 compounds that passed our filters, we found that 5112 (68%) possess one or more stereoisomeric pairs. Notably, 403 of these compounds (7.9%) belong to groups of stereoisomers with variations in BBBP classes, indicating the potential value of incorporating a molecule’s absolute stereochemistry in BBBP prediction models for drug discovery programs.

The structural diversity of these compounds is considerable **(Figure S1a)**, with most belonging to groups of two or more stereoisomers that typically share the same BBBP class **(Figure S1b)**. The size of these stereoisomer groups ranges from 2 to 9 compounds. Interestingly, in the ‘multiclass’ stereoisomer groups—those with varying BBBP classifications—the majority display an equal distribution of BBB+ and BBB-**(Figure S1b, inset)**.

To rigorously evaluate our models, we created subsets in our test sets devoid of stereoisomeric pairs found in the respective training or validation sets of that cross-validation fold, unless the group exhibited variability in BBBP (“Stereoisomers Filtered”). Furthermore, we focused on those molecules in groups of two or more stereoisomers that exhibit variability in BBBP (“Multiclass”) for a more stringent assessment of the model’s discriminative power **(Figure S1c-d)**.

This analysis was extended to stereoisomers and racemates with available log*BB* measurements, some of which showed significant variability **(Figure S1e-f)**. This variation underscores the potential utility of a model capable of discerning how absolute stereochemistry influences BBBP. It should be noted, however, that log*BB* alone does not generally suffice to determine whether stereoisomers differ in free fraction available for efficacy. K_p,uu_ has proven, including in its seminal publication^22^, that stereoselective plasma protein and brain tissue binding can result in different total but the same unbound brain to plasma ratios. Nonetheless, we emphasize the importance of considering absolute stereochemistry from the earliest stages of drug discovery, both in training and evaluating predictive models.

### 3.3. Cross-Validation Results

Our evaluation of various architectures on the full test sets revealed that MPO was significantly outperformed by multiple methods (**Figure S2**, **Table S1)**. The XGBC trained with Mordred descriptors emerged as the top performer, followed closely by Uni-Mol and CANDID-CNS™. For a stricter comparison of our models with MPO, we explored the models’ ability to detect patterns in BBBP by excluding non-multiclass stereoisomers with counterparts in the training or validation sets. This analysis was also aimed to assess the generalizability of ML and NN models compared to MPO.

Interestingly, all models, including MPO, demonstrated reduced performance metrics on stereoisomers filtered test subsets (**Figure 1**, **Table 1**). MPO’s performance showed a decrease of approximately 5% in AUROC on these subsets compared to the full test sets. CANDID-CNS™’s performance decrease is 12%, yet still outperformed MPO by 17-19% according to this metric. This decline in MPO performance could be attributed to a shift towards a more balanced dataset, with fewer disproportionate numbers of BBB+ compared to BBB-compounds. Conversely, the diminished performance of other models on these subsets might indicate insufficient generalization. We considered whether MPO generalizes more effectively than ML models, a hypothesis we will explore further with our internal data in a subsequent section and **(Figure 4)**.

Additionally, we observed that models tend to overestimate performance in scaffold splits, which are typically used for practical absorption, distribution, metabolism, excretion, and toxicity (ADMET) prediction. For example, Uni-Mol and ImageMol models showed lower performance in scaffold splits compared to stratified splits on the full test set, yet outperformed their stratified split versions on the stereoisomers filtered subsets. This discrepancy is likely due to the scaffold split incorporating molecular chirality, leading to stereoisomers of the same group and class appearing in both training and test sets. A scaffold split without chirality consideration might yield results similar to those observed in our stereoisomers filtered test subsets, but potentially with less diverse chemical space coverage in the training set, impacting the model’s generalization ability for internal compounds.

Given these findings, we regard the results from the stereoisomers filtered subsets as a more accurate reflection of potential model performance. In this context, Uni-Mol on a stratified split demonstrated the best performance in key metrics (sensitivity, AUROC, MCC, and accuracy), closely followed by CANDID-CNS™, which tied with Uni-Mol in AUROC (83%) and MCC (0.53) (**Figure 1**, **Table 1**). Notably, both the SVC and the LR trained on calculated MPO scoring function values, such as TPSA, surpassed MPO in AUROC and MCC. We particularly focused on the LR for our comparative analysis with MPO due to its interpretable coefficients.

We observed a similar trend in performance of these models when evaluated on stereoisomer filtered subsets of K_p,uu_ data (**Figure S1**). This suggests that K_p,uu_ and the BBB+/- labels represent similar tasks, as might be expected since K_p,uu_ is the gold standard for quantifying free drug available in the brain. The slight difference in performance may reflect how CNS therapeutics can have low brain penetration but may still be effective due to high potency for example. Alternatively, compounds with non-CNS sites of action may have eluded BBB+ classification in the primary data curation sources^27–29^, even if they have high K_p,uu_, due to a lack of CNS effects. In any case, CANDID-CNS™ trained our processed and filtered version of Meng et al.’s B3DB dataset performs best on K_p,uu_ (**Figure S1**) much as it does here on stereoisomer filtered subsets of the test BBBP data.

Here, we examine the performance of various models on multiclass stereoisomer subsets of the test sets (**Figure 2**, **Table 2**). Among all models, the MolCLR GIN showed the highest MCC (0.37±0.15), albeit with a large standard deviation, while CANDID-CNS™’s AUROC (0.62±0.05) was two standard deviations above random. MPO’s AUROC was comparable to a random classifier on these subsets. This outcome was anticipated given MPO’s general lack of consideration for stereoisomerism. The sample size here is small (20-50 compounds), but the superior performance of AttFP GNN models including CANDID-CNS™ is promising. However, Uni-Mol exhibited inferior performance relative to the MolCLR and AttFP models on multiclass subsets. The superior performance of the AttFP models raise questions as to whether the pretraining methods used in MolCLR, Uni-Mol, and ImageMol prepare them for finetuning on the BBBP task. Further investigation is warranted to determine if factors like pretraining hyperparameters contribute to these results. An intriguing aspect for future exploration is the representation of stereoisomers in the pretraining sets of these models.

While these results do not conclusively indicate an exceptional ability of MolCLR GIN and AttFP models to classify multiclass stereoisomers in BBBP, they suggest potential if supplemented with more labeled data. This inference is supported by the fact that CANDID-CNS™’s MCC (0.34±0.09) on multiclass test subsets compares favorably with MPO’s MCC (0.29±0.04) on stereoisomers filtered test subsets. Graph representations, which incorporate atom and bond stereoisomerism, along with attention, likely contributed to learning patterns related to BBBP and 3D structure, as evidenced by these results.

Given that multiclass performance might underestimate the utility of these models, we view it instead as an indication of their ability to discern minor structural changes that significantly affect molecular properties, akin to activity cliffs^67^. Therefore, we focus CANDID-CNS™ in subsequent sections for a more detailed comparison with MPO, probing their capabilities in depth. In our efforts to enhance model performance for stereoisomers, we incorporated recent advancements from Gaiński et al., specifically their GNN architecture emphasizing chirality^68^, into our AttFP GNN model by adding a chiENN layer. This integration resulted in a 3% increase in the AttFP GNN’s AUROC, though it also led to a rise in standard deviation from 0.05 to 0.10.

### 3.4. Detailed Comparison of CANDID-CNS™ with MPO on a Stereoisomers Filtered Test Subset

Here, we compared the performance of MPO and CANDID-CNS™ on a stratified test set, where stereoisomers were selectively removed based on their group’s inclusion in the training and validation sets, while retaining all multiclass stereoisomers. This approach was designed to ascertain the strengths and limitations of each method. This difference is illustrated in **Figure 3a** with a decision boundary scatterplot and the corresponding regional confusion matrices **(Figure 3b-e)**. The optimal decision boundaries, aiming to maximize the difference between the true positive rate and the false positive rate, are highlighted. We further analyzed specific structures within this test subset to understand the compatibility and unique attributes of both approaches.

**Figure 3.**
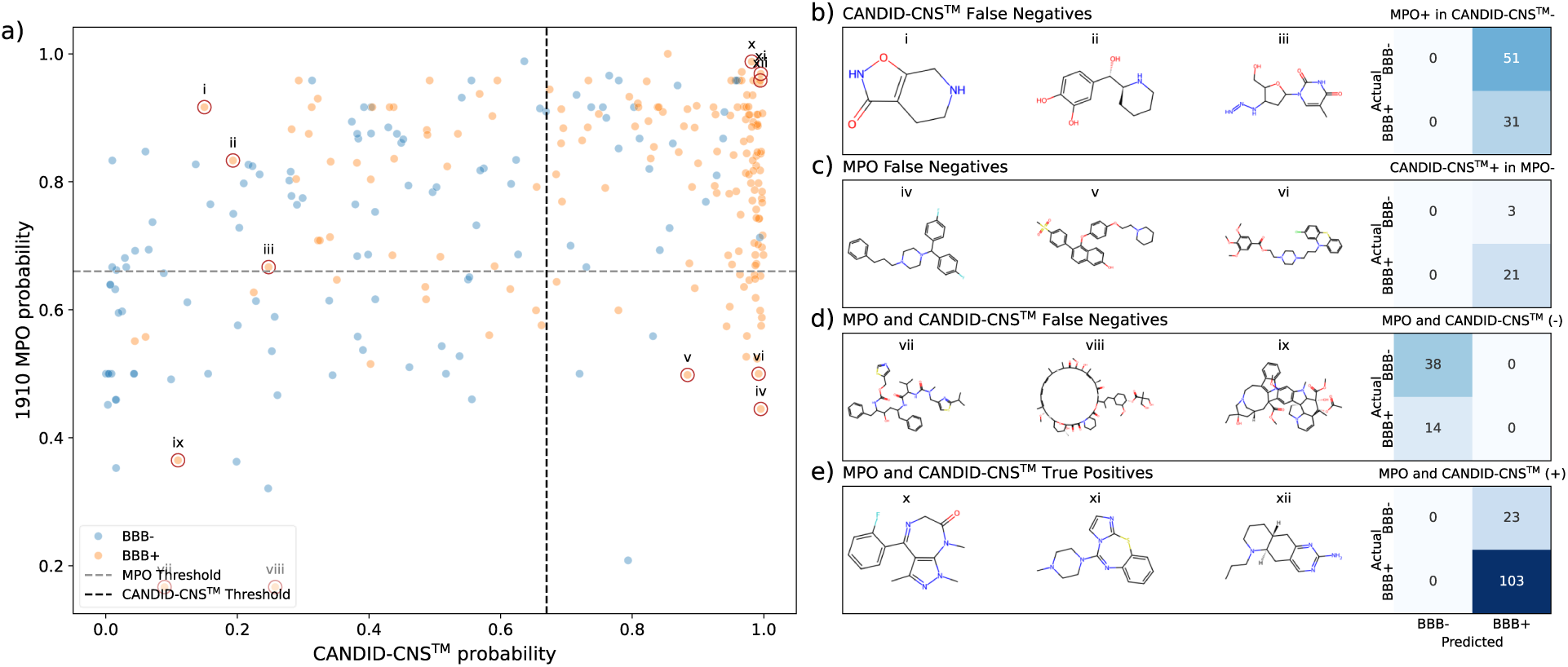
CANDID-CNS™ outperforms MPO on a stereoisomer filtered test subset. a) Decision boundaries between CANDID-CNS™ and MPO. b) CANDID-CNS™ false negatives (i-iii) and partial confusion matrix for this upper left region in (a), showing MPO predicts 51 false positives. c) MPO false negatives (iv-vi) and partial confusion matrix for the lower right region in (a), showing CANDID-CNS™ predicts only 3 false positives here. d) Lower left region, where MPO and CANDID-CNS™ both classify 14 false negatives, including molecules vii-ix, shown. e) Upper right region, where MPO and CANDID-CNS™ select 103 true positives, including molecules x-xii, shown, and 23 false negatives.

In addition to better AUROC and MCC metrics, the CANDID-CNS™’s AUPRC is higher by about 0.15 compared to MPO. The dense scatter of positive orange points at the right of **Figure 3a** is related to this statistic. At the top of the plot, however, where MPO is highest, orange points do not appear to cluster much more densely than blue negatives. In terms of compound prioritization, choosing the highest probability compounds from the CANDID-CNS™ would result in more true positives than choosing from the highest MPO scoring compounds in this test set.

In the upper left quadrant of **Figure 3a**, MPO identifies additional BBB+ compounds compared to CANDID-CNS™ but also includes more false positives. Conversely, in the lower right quadrant, CANDID-CNS™ identifies true positives missed by MPO with fewer false positives.

Analysis of CANDID-CNS™ false negatives (upper left, **Figure 3a**, i-iii) shows compounds that are smaller than the MPO false negatives (iv-vi) illustrated from the lower right quadrant (iv-vi) which the CANDID-CNS™ identifies as true positives. (i-iii) are relatively small compounds that follow MPO guidelines of modest Mw, logD/P, TPSA and pKa. The fact that CANDID-CNS™ does not classify these correctly suggests future avenues for improvement.

On the other hand, CANDID-CNS™ correctly classifies compounds such as (iv-vi) that are likely too large and hydrophobic to score well with MPO. (iv-vi) have lipophilic alkyl chains and 3-4 aromatic rings each, implying they have high property forecast index (PFI)^69^. However, the alkyl chains in (iv-vi) are flexible. Flexibility is a property that may contribute to their permeability, and a pattern which CANDID-CNS™ may have learned. Molecule (iv) has two fluorinated carbons, which has been increasingly found in FDA approved small molecule therapeutics^70^. It is believed that fluorine may aid in permeability, as well as having other beneficial effects for drug binding and metabolic stability^71^. Visualization of bond attention weights suggests the fluorine atoms and sp3 carbon chains were learned features underlying CANDID-CNS™’s correct classification of compound (iv) **(Figure S8)**.

Molecules in (**Figure 3d**, vii-ix) suggest areas for further improvement to both CANDID-CNS™ and MPO. These are very large molecules including a macrocycle (viii). (iv) is another flexible molecule with many sp3 carbons, but CANDID-CNS™ did not label it positive. (ix) may not have as many sp3 carbons, and thus may not be as flexible, as either (vii) or (viii), but it likely has a complex 3D structure that may be important for its permeability.

### 3.5. Comparison of BBBP GNNs with MPO on Internal Data

Here we discuss the relevance of the processed dataset and our models for predicting BBBP retrospectively on a small set of internal compounds based on a common scaffold. Internal and 3^rd^ party MPO scores (ACD Percepta, Advanced Chemistry Development, Inc., Toronto, Ontario) were used prospectively as part of the prioritization of these compounds, so this is not a random selection.

Figure 4a presents a t-SNE plot of principle component analysis (PCA) reduced ECFPs of 29 internal compounds alongside the preprocessed BBBP dataset. This visualization indicated that our compounds were not markedly dissimilar from the broader dataset in the high dimensional ECFP space, as they clustered within the main body of the dataset. Despite varying log*BB* values, these compounds also showed a tendency to cluster together, highlighting their similarity in ECFP representation. Further analysis involved identifying the most similar molecules in the processed dataset for each of the 29 compounds (Figure 4b). For each compound, we quantified the number of molecules in the dataset exceeding a 0.27 Tanimoto similarity threshold, which is considered significant for predictive accuracy^72,73^, barring potential activity cliffs^67^.

**Figure 4.**
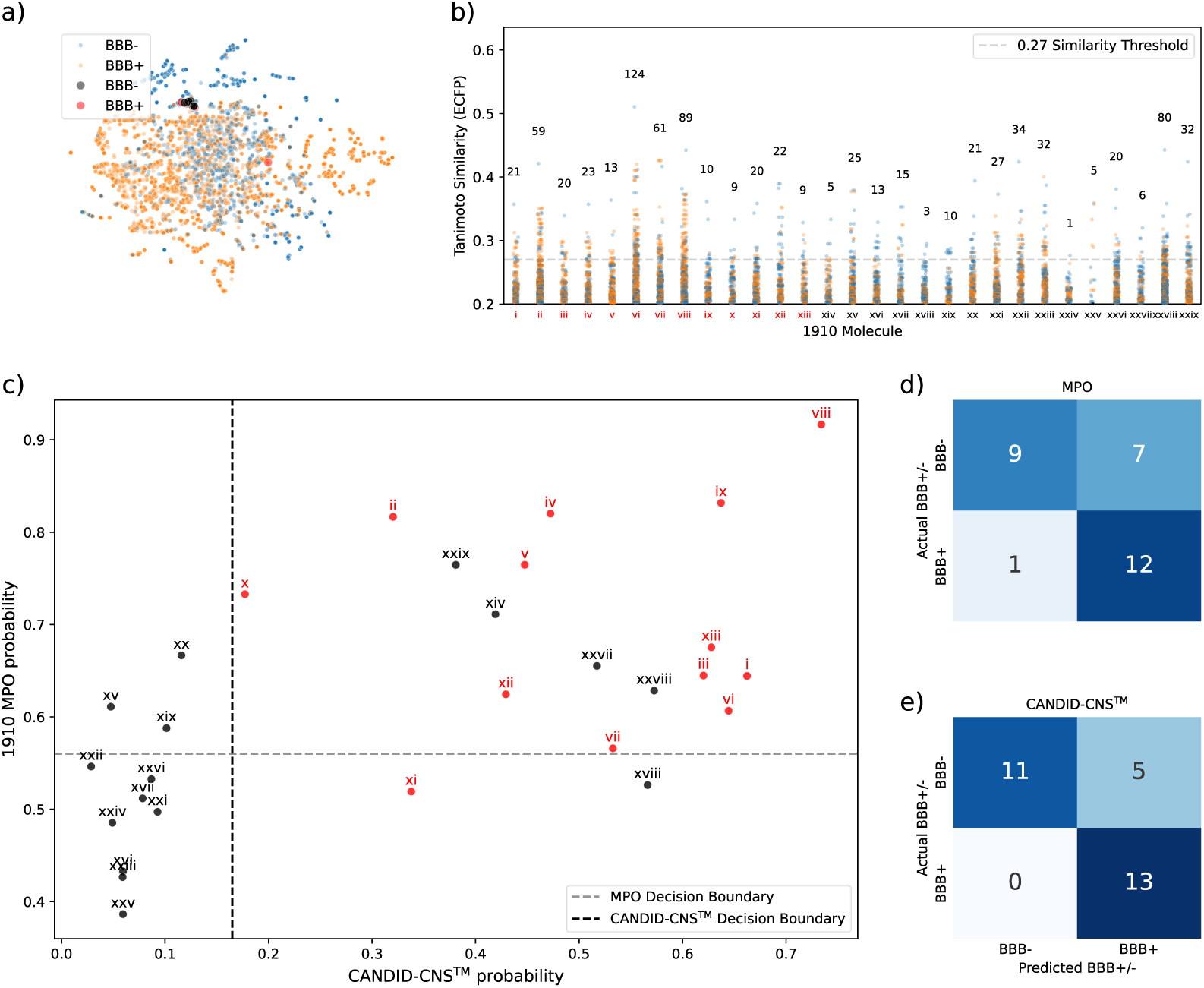
CANDID-CNS™ outperforms MPO on internal data. a) t-SNE showing most internal compounds cluster in one region among BBB+ and BBB-compounds. b) Strip plot showing the similarity of the most similar molecules in the processed dataset to each internal compound, with counts of molecules >0.27 similarity shown above each strip. Compounds are numbered in roman numerals and ordered from highest log*BB* value (i) to lowest (xxix). c) Decision boundaries of 1910 MPO and CANDID-CNS™ on internal compounds. (d-e) Confusion matrices of data in (c) showing CANDID-CNS™ outperforms MPO.

Despite the range of the most similar molecules for each compound being quite broad (from 1 to 124), we argue that this diversity was sufficient for training models capable of accurately predicting the BBBP of our internal compounds. The comparative performance of CANDID-CNS™ and MPO on these internal compounds is depicted in Figure 4c, with their respective confusion matrices shown in Figure 4, subpanels d and e. Interestingly, both MPO and CANDID-CNS™ showed comparable efficacy on this small evaluation set, with the GNN slightly outperforming MPO. The GNN’s optimal threshold identified an additional true positive compared to MPO, avoiding any false negatives and registering two fewer false positives.

Our findings indicate that CANDID-CNS™ was equal to or surpassed MPO in terms of overall classifier metrics such as AUROC, MCC, and accuracy. MolCLR also showed a comparable performance. However, the most effective models on this dataset were the SVC and LR models trained using TPSA, MW, clog*P*, clog*D*, most basic pKa, and HBD as features. These results are summarized in **Table 3**.

**Table 3.** MPO and ML Model Comparison on Internal Data.

| Classifier | AUROC | Sensitivity | MCC | Accuracy | Precision | Specificity |
| --- | --- | --- | --- | --- | --- | --- |
| 1910 MPO | 0.81 | 0.92 | 0.51 | 0.72 | 0.63 | 0.56 |
| CDD MPO | 0.90 | 0.77 | 0.72 | 0.86 | 0.91 | **0.94** |
| CANDID-CNS™ (Stratified) | 0.90±0.01 | 0.97±0.06 | 0.80±0.04 | 0.89±0.03 | 0.83±0.06 | 0.82±0.07 |
| MolCLR GIN (Stratified) | 0.81±0.05 | 0.82±0.08 | 0.63±0.12 | 0.81±0.06 | 0.78±0.08 | 0.81±0.07 |
| LR MPO (Stratified) | 0.96±0.00 | **1.00±0.00** | **0.87±0.00** | **0.93±0.00** | **0.87±0.00** | **0.88±0.00** |
| SVC MPO (Stratified) | **0.98±0.00** | **1.00±0.00** | **0.87±0.00** | **0.93±0.00** | **0.87±0.00** | **0.88±0.00** |
| TPSA score | 0.95 | **1.00** | **0.87** | **0.93** | **0.87** | **0.88** |

## 4. Conclusion

In conclusion, our findings indicate that the MPO approach can be improved based on the insights gained from our study. We propose a recalibration of the MPO scoring function, suggesting a MW threshold at 500 Da instead of the conventional 400 Da. This recommendation is grounded in the patterns observed in the histograms presented in **Figure S4a**. Additionally, we emphasize the predictive capacity of the TPSA in determining BBBP. Our analysis supports the notion of assigning a greater weight to TPSA in MPO calculations, such as its utilization in our LR models. Furthermore, the data **(Figure S7)** reveal the potential of Chromlog*D* in distinguishing BBB+ from BBB-compounds. Intriguingly, Chromlog*D*, which is influenced by molecular volume, holds promise for differentiating stereoisomers. However, it is crucial to acknowledge that while our analysis suggests that a relaxation of MPO’s lipophilicity constraints might enhance BBBP prediction, these parameters remain vital for addressing a spectrum of other ADMET and toxicity outcomes, notably hERG inhibition liability^25,45^. Therefore, any modifications to the MPO framework must carefully balance the improved prediction of BBBP with the preservation of its applicability to other critical ADMET and toxicity endpoints.

Although MPO has limitations in BBBP prediction, it’s crucial to remember its broader goal of identifying compounds with generally favorable drug properties. The original MPO framework by Wager et al.^25,45^ referenced Lipinski’s rules^74^ and considered studies linking clog*P* and TPSA with toxicity^75^, emphasizing the inclusion of lipophilicity and TPSA not for BBBP prediction accuracy, but to minimize potential off target interactions. In this context, using a lipophilicity classifier or regressor, alongside a BBBP classifier, could serve as a modified MPO approach, balancing BBBP prediction with general drug likeness and off target minimization. Furthermore, integrating GNN models for direct toxicity prediction^76^ with a GNN BBBP classifier could facilitate simultaneous prioritization of both endpoints.

Because of the importance of TPSA in BBBP prediction, it may be interesting to study alternative methods of calculating PSA. One avenue of study is whether quantum chemical calculations^77^ and vectors of 3D PSA values for multiple conformers could improve BBBP classification. Experimental polar surface area (EPSA)^78^ measurements may also be useful for improving upon TPSA and thus, perhaps, BBBP prediction.

Regarding the cross-validation performance of the state of the art NN and ML methods, the emergence of CANDID-CNS™ as the top performer in the increasingly strict stereoisomer based test subsets carries certain important implications. First, higher parameter models and pretraining strategies did not yield better performance in the cases of Uni-Mol and ImageMol. Similarly, MolCLR’s contrastive learning pretraining did not compensate for its lack of an attention mechanism when compared to CANDID-CNS™. These results suggest that the attention mechanism is more important for BBBP prediction than the various pretraining strategies employed by these other approaches.

The surprising performance of CANDID-CNS™ on the strictest test subset of multiclass stereoisomers is reflected in its AUROC and MCC rivaling or outperforming MPO’s AUROC and MCC on the less strict stereoisomers filtered subset. CANDID-CNS™’s performance overall and on this strict test case suggest that explicit 3D positional embeddings, such as those Uni-Mol and the ChiENN Attentive GNN implementation use, may not be as useful for stereochemistry-related BBBP prediction as stereochemical atom (node) and bond (edge) graph encodings. Ultimately, more studies would be required to determine if, for example, explicit 3D embeddings via quantum mechanical calculations or molecular dynamic simulations could improve these features for BBBP prediction.

Finally, we also note an avenue for future investigation of CANDID-CNS™ and its comparison with descriptor models. Descriptor methods are useful because they provide human interpretable features. However, both the XGBC and SVC descriptor models perform very poorly on the strictest test subset of multiclass stereoisomers. An attention coefficient analysis could reveal which features CANDID-CNS™ determines are most responsible for stereoisomeric differences in BBBP. Further analysis of such features could point to new descriptors relevant to BBBP and stereoselectivity.

The ultimate usefulness of MPO, ML, and NN classifiers for BBBP will be most effectively established through extensive comparative studies involving a significant number of designed and prioritized compounds. This validation should encompass a broad range of assessments, including physical chemistry analyses, ADMET profiles, *in vivo* PK, toxicity evaluations, drug efficacy trials, and clinical outcomes. It is also noteworthy that enhancing potency and selectivity for specific targets may reduce the likelihood of adverse effects by decreasing the required dosage^75^, potentially favoring the application of BBBP models over traditional MPO methods in early drug development phases, such as hit to lead prioritization or the optimization of nano or picomolar binders. The continued exploration and comparison of ML, NN, and MPO methodologies in the development of CNS drug candidates are imperative, with an emphasis on both BBBP prediction and the integration of broader ADMET considerations and drug efficacy.

## ASSOCIATED CONTENT

The document SI.pdf contains Supplementary Information.

## AUTHOR INFORMATION

### Corresponding Author

Jennifer O. Nwankwo

1910 Genetics, 451 D Street, Suite 905, Boston, MA, 02210

### Author contributions

JON conceptualized the original study and drug discovery program. JON, BM, and JWC planned and coordinated the BBBP ML research. JD and JWC analyzed the dataset, plotted similarity scores, and compared public with internal data. ME, MA, JWC, JD, MG, and DR coordinated the preprocessing of the data. DR preprocessed and split the data. ME, DR, and JWC wrote cross-validation and ROC curve fitting code. JWC performed the stereoisomer analysis and wrote stereoisomer filtering codes. MA and LPV advised on the filtering of charged compounds. LPV designed and implemented the ChromLog*D* method, advised on chemical structures, and provided literature references. MG implemented Uni-Mol with ME and DR. DR optimized hyperparameters for the XGBC. JWC optimized hyperparameters for the SVC. DR cross-validated Uni-Mol, ImageMol, MolCLR and the XGBC. JWC and JD cross-validated the SVCs. JWC cross-validated the AttFP GNNs, LR, and calculated MPO scores. JWC plotted figures. JWC and ME wrote the manuscript. JON, DR, BC, BM, LPV, and MA reviewed and contributed to the manuscript.

## FUNDING SOURCES

This work was funded by investors in 1910 Genetics as well as a Small Business Technology Transfer (STTR) grant (Award Number R41NS118992-01) to J.O.N from the Helping to End Addiction Long-term (HEAL) initiative at the National Institute of Neurological Disorders and Stroke (NINDS) of the National Institutes of Health (NIH).

## ACKNOWLEDGMENT

We acknowledge Mark Ashwell for providing a background reference on brain and CNS penetration of small molecules. We acknowledge Bernardo Hernandez-Adame for helpful discussions and Bernardo Hernandez-Adame and Shannon Cosgrove for initial implementations of our MPO calculator. We acknowledge Peter Bertinato for helpful discussions and anonymous reviewers for their feedback.

## ABBREVIATIONS

ML: machine learning
TPSA: topological polar surface area
BBBP: blood-brain barrier permeability
MW: molecular weight
HBD: hydrogen bond donors
log*P*: log of partition coefficient of neutral species
log*D*: log of molecular distribution (e.g. water/octanol partitioning)
Chromlog*D*: chromatographic log*D*
AUROC: area under receiving operating characteristic curve
MCC: Matthews Correlation Coefficient
CNS: central nervous system

## DATA AND SOFTWARE AVAILABILITY

Our curated splits of the B3DB dataset into training, validation, and test sets are available. We also provide the stereoisomers filtered and multiclass subsets of the test sets. All ML and NN model architectures are available with hyperparameters and training procedures described in detail for reproducibility.

Internal compound structures are proprietary and cannot be made available at this time pending intellectual property protection. Our essential findings including the analysis of MPO and its outperformance by ML and NN models do not require internal data for reproduction.

## COMPETING INTERESTS STATEMENT

All authors are employees, contractors, and/or shareholders of 1910 Genetics.

## Notes

### Competing Interest Statement

All authors are or have been employees, contractors, and/or shareholders of 1910 Genetics.

### Summary of Updates

Moved figures 1 and 2 to supplement Figure references updated Minor text changes

