## Supplementary Information for "Attentive graph neural network models for the prediction of blood-brain barrier permeability"

**Supplementary Information: Detailed Data Curation Process for the B3DB Dataset**

Our data curation process for the B3DB dataset was designed to ensure the quality and relevance of the data for our ML analyses. The initial dataset comprised 7809 compounds. We began by removing two compounds with missing SMILES (Simplified Molecular Input Line Entry System) strings and two compounds where SMILES sequences were incorrectly entered as numeric strings. This refinement reduced the dataset to 7805 compounds.

We then focused on the chemical properties of the compounds. Our analysis revealed that 324 compounds possessed a net charge, and approximately 173 compounds, while net neutral, had multiple charged atoms. Among these, some molecules exhibited positively charged carbons, an indication of incorrect SMILES strings. Additionally, certain molecules, such as sulfonium and ammonium cations, were removed because they did not align with the focus of our drug development interests. This charge-based filtering led to the exclusion of these 497 compounds.

The next step in our process was to preprocess the molecular structures for ML compatibility. We selected only the freebase form of each molecule, which entailed keeping the largest fragment in cases where a molecule consisted of multiple fragments. We also standardized the structures through protonation, preparing them for 3D embedding, a requirement for models like Uni-Mol and chiENN[1]. During this phase, we identified three pairs of duplicate molecules. Upon inspection, we decided to retain ampicillin, probenecid, and the first listed molecule of colestipol, thus removing the duplicates.

Finally, we subjected the molecules to the Uni-Mol preprocessing script to generate 3D conformations. During this stage, 13 molecules consistently failed to embed properly in 10 separate attempts. To ensure uniformity in featurization methods and architectural comparisons, these compounds were removed from the dataset, resulting in the final dataset of 7465 molecules.

**Supplementary Information: Detailed Architecture Hyperparameters and Training**

In the process of training our machine learning models, we undertook an extensive approach to optimize each architecture's performance. For the SVC implementation using scikit-learn, we conducted a grid search over hyperparameters on a subset of Roy et al.[27] BBBP dataset, prioritizing specificity in our scoring. We finalized the SVC with a regularization parameter of 100 and a Gaussian radial basis function kernel with a coefficient of 0.5.

For XGBoost, the hyperparameters selected via grid search included a learning rate of 0.05, a maximum depth of 20, and 200 estimators. The model's gamma, alpha, and lambda were set to 0, with a scale position weight of 5 and subsample of 0.9.

Our CANDID-CNS™ adaptation of the AttFP GNN included a sigmoid layer for classification, with 39 input channels, 200 hidden channels, and a dropout of 0.2. The atom features included atomic number, degree, hybridization, aromaticity, hydrogens, and chirality. The bond features represented 10 edge dimensions. The optimizer used was Adam with a learning rate of $10^{-2.5}$ and weight decay of $10^{5}$.

For the chiENN version of AttFP GNN, we integrated a chiENN layer and modified hyperparameters to accommodate chiENN edge graphs, including increasing the number of input channels to 88 and the edge dimension to 59.

The pre-trained ImageMol model was fine-tuned with a binary cross-entropy with logits loss function, a learning rate of 0.01, and an early stopping patience of 30 epochs. MolCLR models were fine-tuned over 100 epochs with a learning rate of $5\times10^{-5}$ and a weight decay of $10^{-6}$. The models had 5 layers, an embedding dimension of 300, and an output feature dimension of 512.

For Uni-Mol, we used the pre-trained model weights and adhered to the default training script, including a dropout of 0, a warmup parameter of 0.06, and a learning rate for the Adam optimizer set to $10^{-4}$.

**Supplementary Information: Detailed Definitions and Equations for Evaluation Metrics**

In our study, we employed a range of standard classifier evaluation metrics. Here, we provide the detailed equations and definitions for each:

1. **Sensitivity**: Also known as the true positive rate, sensitivity measures the proportion of actual positives correctly identified by the classifier. It is calculated as:

$$sensitivity = \frac{TP}{TP + FN}$$

where TP is the number of true positives and FN is the number of false negatives.

2. **Accuracy**: Accuracy measures the proportion of correct predictions (both true positives and true negatives) among the total number of cases examined. It is given by:

$$Accuracy = \frac{TP + TN}{TP + TN + FP + FN}$$

where TN is the number of true negatives and FP is the number of false positives.

3. **Precision**: Precision, or positive predictive value, measures the proportion of positive identifications that were correct. It is calculated as:

$$Precision = \frac{TP}{TP + FP}$$

4. **Specificity**: Also known as the true negative rate, specificity measures the proportion of actual negatives correctly identified. It is defined as:

$$Specificity = \frac{TN}{TN + FP}$$

5. **Receiver Operating Characteristic (ROC) Area Under Curve (AUC)**: This metric evaluates the classifier's ability to distinguish between classes. The ROC curve plots the true positive rate against the false positive rate at various threshold settings.

6. **Matthews Correlation Coefficient (MCC)**: MCC is a statistical rate which produces a high score only if the prediction obtained good results in all of the four confusion matrix categories (TP, FN, TN, and FP), proportionally both to the size of positive elements and the size of negative elements in the dataset. The MCC is calculated as:

$$MCC = \frac{(TP \times TN) - (FP \times FN)}{\sqrt{(TP + FP)(TP + FN)(TN + FP)(TN + FN)}}$$

**Supplementary Information: Detailed Lipophilicity Measurement Methods and Model Training**

In developing our computational models for lipophilicity, we used two AttFP GNN-based models for log*D* and Chromlog*D* calculations. The log*D* model's training incorporated 1.4 million ChemAxon clog*D* values from ChEMBL[56], along with additional measurements from Analiza and Aurigene Services Limited. Its hyperparameters were the same as CANDID-CNS™ model, except for a learning rate of $10^{-4}$, with training manually stopped after around 500 epochs upon observing an increase in validation loss.

The Chromlog*D* model was trained using a slightly modified AttFP model, incorporating a node feature size of 74 and an edge feature size of 12. The graph feature size was set at 256, with a dropout of 0.2. We employed a learning rate of $10^{-3}$ and an early stopping mechanism with a patience of 20 epochs. The Adam optimizer and negative root mean squared error were used as the loss function. The clog*D* ChemAxon values were rescaled to align with the ChromLogD scale using a set of 120 dual log*D*/Chromlog*D* measurements and a linear transformation derived from this dataset.

**Supplementary Information: B3DB Analysis and Stereoisomer Test Subsets**

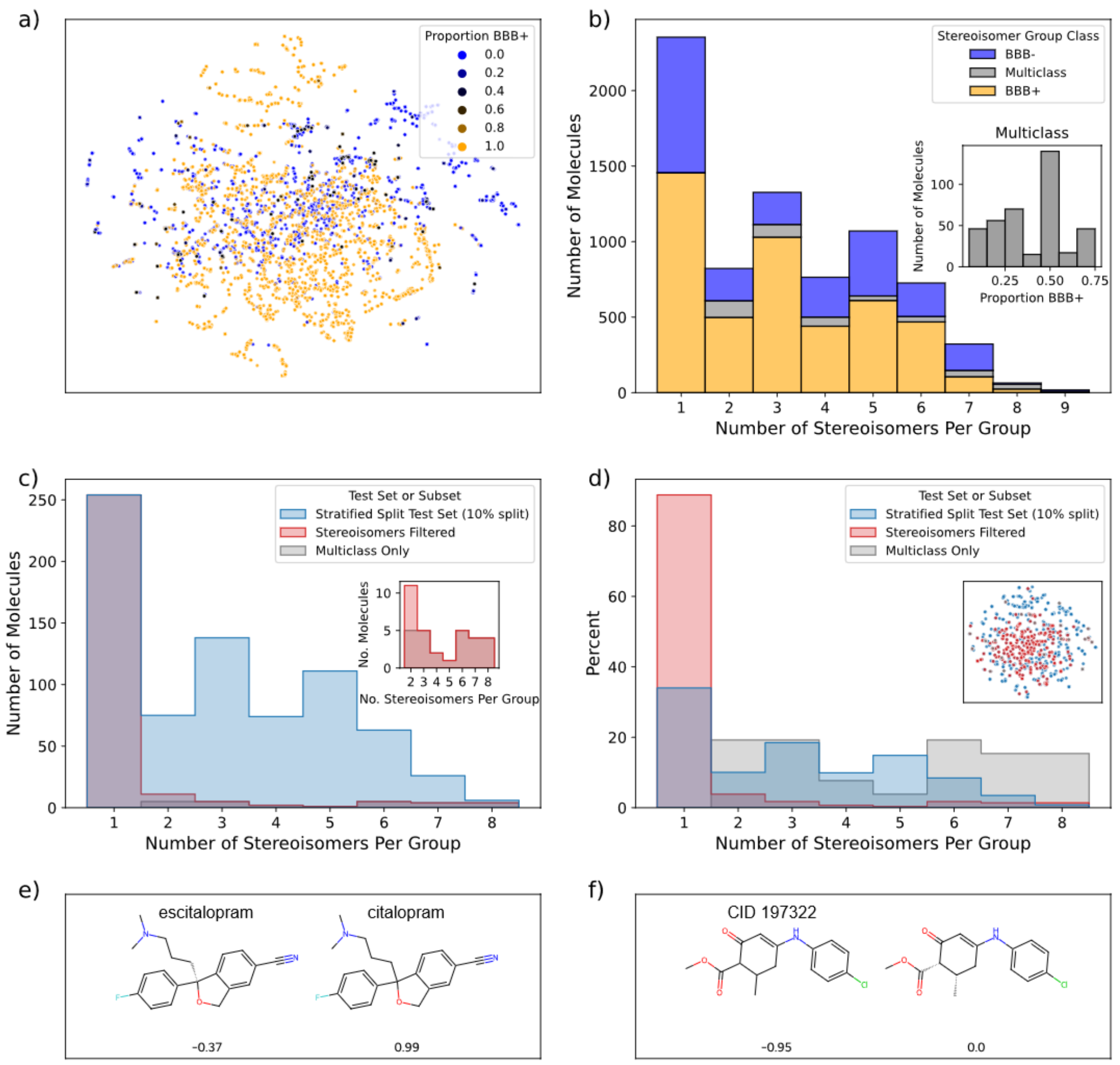


**Figure S1.** Prevalence and importance of stereoisomerism in BBBP Data. (a) t-distributed stochastic neighbor embedding (t-SNE) plot based on Extended Connectivity Fingerprints (ECFPs) showing the variety of all structures and the proportion BBB+ of their stereoisomer groups (0, 1, or intermediate for multiclass). (b) The distribution of the data by the number of stereoisomers in each group. 1 indicates a unique molecule without a stereoisomeric pair in the dataset. Inset: the number of compounds in multiclass stereoisomer groups by proportion BBB+. (c-d) Example test set and its stereoisomers filtered and multiclass subsets. (c) Distribution by number of molecules. Inset: focus on the non-unique (N>2) stereoisomer groups in the stereoisomers filtered and multiclass subsets. (d) Distributions in percent, showing the stereoisomers filtered data is mostly unique molecules without a stereoisomeric pair. Inset: t-SNE of this test set colored by the smallest subset in which the molecule belongs (full test set, stereoisomers filtered, or multiclass). (e-f) Two pairs of racemates and corresponding stereoisomers that maximize the difference in reported logBB measurements (shown beneath the molecules). (e) escitalopram and its higher logBB racemate. (f) A low logBB racemate and its higher logBB stereoisomer from anticonvulsant studies[66].

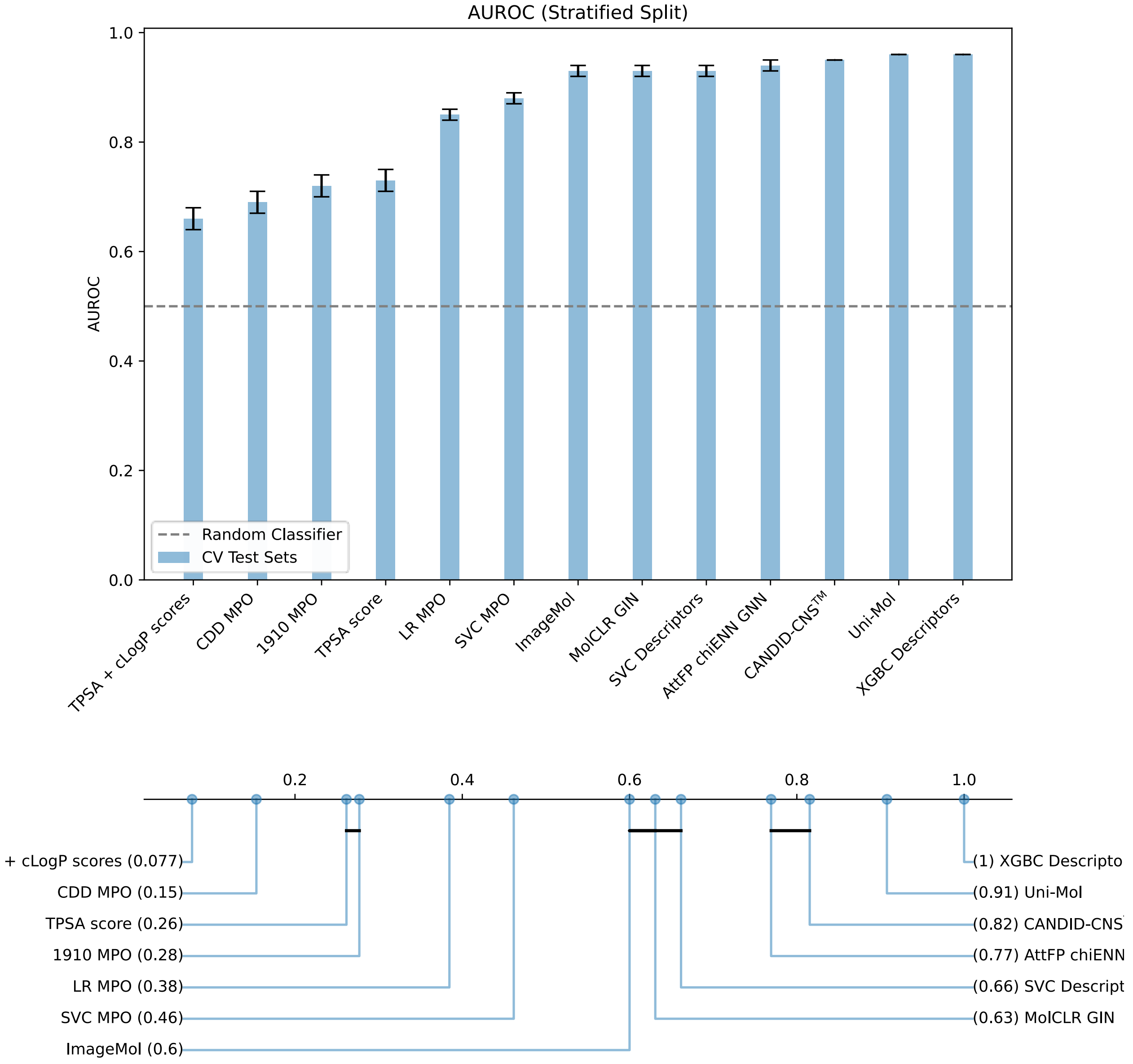


**Figure S2.** ML/NN Models Outperform MPO on Cross-Validation Test Sets. a) Models ordered by AUROC cross-validation average. Error bars represent one standard deviation from the mean. b) Critical difference diagram showing which models are statistically similar by Friedman rank. Models are connected if pairwise Conover p-value is > 0.05.

**Table S1.** Model Comparison on Cross-Validation Test Sets. Data splitting is stratified unless otherwise noted.

| Classifier | AUROC | MCC | AUPRC | Sensitivity | Accuracy | Precision | Specificity |
|---|---|---|---|---|---|---|---|
| Uni-Mol | **0.96±0.00** | **0.79±0.01** | **0.97±0.00** | **0.91±0.02** | **0.90±0.01** | 0.94±0.01 | 0.89±0.02 |
| XGBC Descriptors | **0.96±0.00** | **0.79±0.02** | 0.98±0.00 | 0.89±0.04 | **0.90±0.01** | 0.95±0.02 | **0.91±0.03** |
| Uni-Mol (Random) | 0.95±0.01 | 0.76±0.02 | 0.97±0.01 | 0.87±0.03 | 0.88±0.01 | 0.94±0.01 | 0.90±0.01 |

| | | | | | | | |
|---|---|---|---|---|---|---|---|
| CANDID-CNS™ | 0.95±0.00 | 0.75±0.02 | 0.97±0.00 | 0.89±0.03 | 0.88±0.01 | 0.93±0.02 | 0.87±0.04 |
| AttFP chiENN GNN | 0.94±0.01 | 0.72±0.03 | 0.96±0.01 | 0.84±0.03 | 0.86±0.02 | 0.94±0.00 | 0.90±0.01 |
| ImageMol | 0.93±0.01 | 0.75±0.01 | 0.95±0.01 | 0.90±0.01 | 0.88±0.01 | 0.91±0.02 | 0.85±0.03 |
| MolCLR GIN | 0.93±0.01 | 0.70±0.01 | 0.95±0.01 | 0.86±0.02 | 0.86±0.01 | 0.92±0.01 | 0.86±0.03 |
| SVC Descriptors | 0.93±0.01 | 0.76±0.02 | 0.95±0.01 | 0.88±0.02 | 0.89±0.01 | 0.94±0.01 | 0.89±0.02 |
| ImageMol (Random) | 0.92±0.00 | 0.69±0.01 | 0.95±0.00 | 0.84±0.02 | 0.85±0.01 | 0.93±0.01 | 0.88±0.02 |
| Uni-Mol (Scaffold) | 0.91 | 0.61 | 0.97±0.00 | 0.86 | 0.85 | 0.95 | 0.83 |
| ImageMol (Scaffold) | 0.90 | 0.56 | 0.97±0.00 | 0.80 | 0.81 | **0.96** | 0.87 |
| SVC MPO | 0.88±0.01 | 0.63±0.03 | 0.91±0.01 | 0.82±0.05 | 0.82±0.02 | 0.89±0.01 | 0.82±0.03 |
| LR MPO | 0.85±0.01 | 0.60±0.02 | 0.88±0.01 | 0.86±0.02 | 0.81±0.01 | 0.85±0.00 | 0.73±0.01 |
| TPSA score | 0.73±0.02 | 0.48±0.03 | 0.77±0.01 | 0.85±0.02 | 0.76±0.01 | 0.80±0.01 | 0.61±0.02 |
| 1910 MPO | 0.72±0.02 | 0.39±0.04 | 0.80±0.02 | 0.75±0.04 | 0.71±0.02 | 0.79±0.02 | 0.64±0.04 |
| CDD MPO | 0.69±0.02 | 0.32±0.04 | 0.76±0.02 | 0.67±0.03 | 0.66±0.01 | 0.76±0.03 | 0.65±0.06 |
| TPSA + cLogP | 0.66±0.02 | 0.38±0.03 | 0.75±0.01 | 0.73±0.01 | 0.70±0.01 | 0.79±0.02 | 0.66±0.03 |

### Supplementary Information: $K_{p,uu}$ Performance

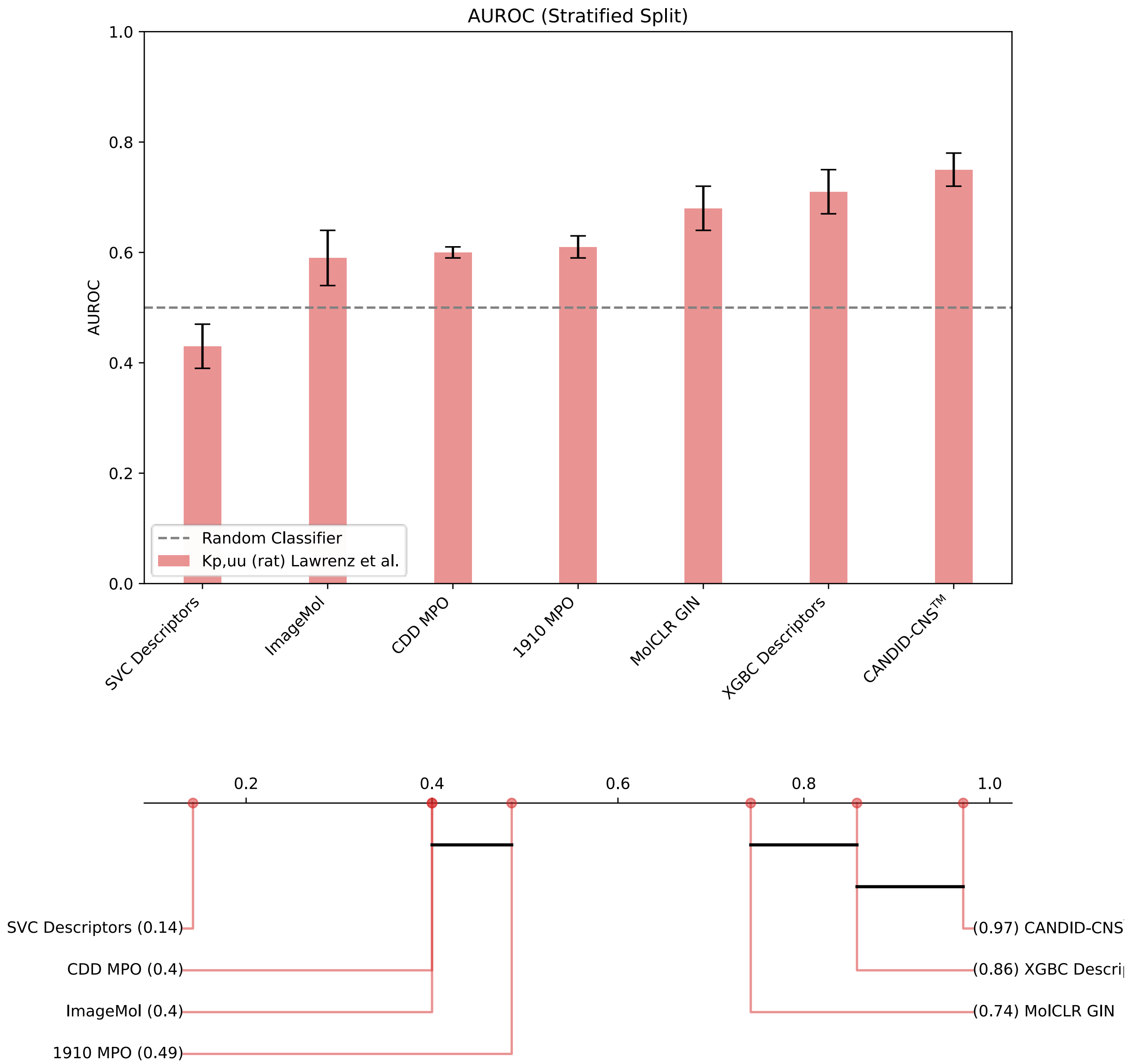


**Figure S3. CANDID-CNS™ Outperforms All Models on $K_{p,uu}$ Data.** CANDID-CNS™ is also the most performant model on stereoisomer filtered test sets of Lawrenz et al.'s $K_{p,uu}$ data and again significantly more performant by critical difference analysis than MPO models. The fact this and other ML and NN models are predictive on $K_{p,uu}$ without being trained with explicit $K_{p,uu}$ data suggest that the BBB+/- class information is closely related to $K_{p,uu}$.

### Supplementary Information: MPO Analysis

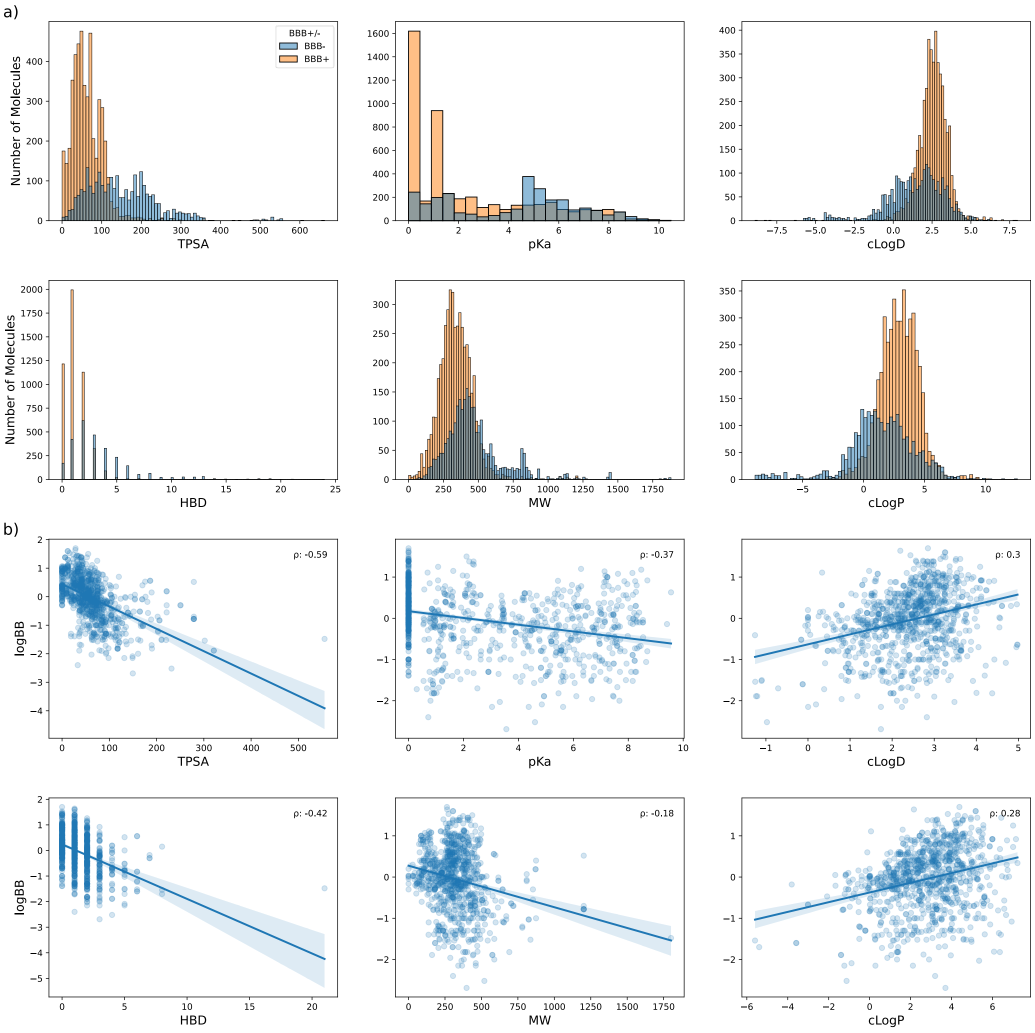


**Figure S4.** Problems with MPO for Predicting BBBP. a) Each MPO component alone does not predict BBBP class. b) Log*BB* is most anticorrelated with TPSA (Spearman ρ = -0.59), suggesting it should be weighted more heavily. clog*P* and clog*D* values are correlated with log*BB*, suggesting that MPO scoring of these properties does not improve its accuracy for predicting CNS penetration.

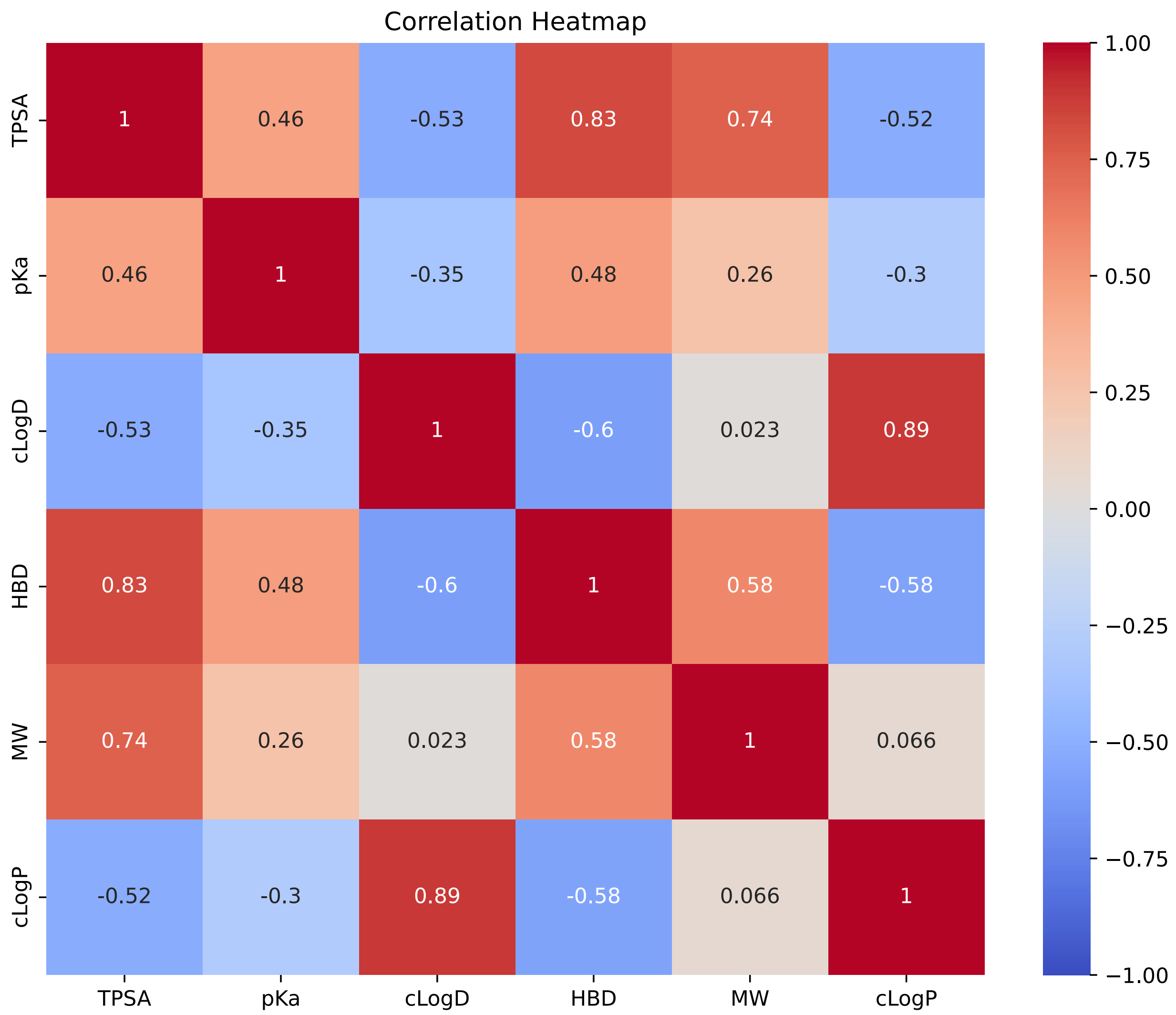


**Figure S5.** Most MPO Properties are Strongly Correlated or Anticorrelated with Each Other. The correlation and anticorrelation suggests a degree of redundancy in the use of these descriptors for predicting BBBP.

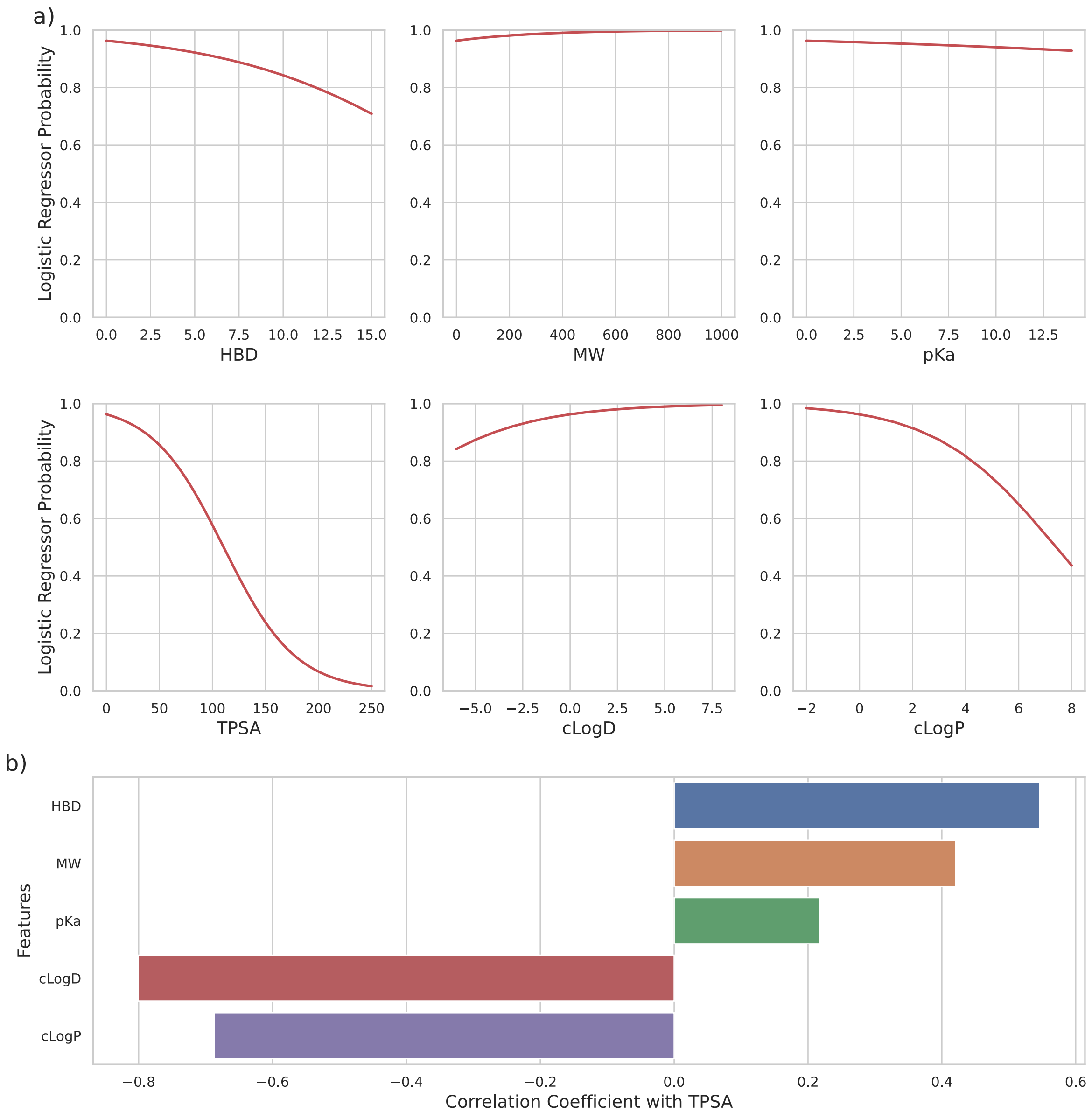


**Figure S6.** Logistic Regression Suggests MPO Components Are Not Weighted Optimally for BBBP Prediction. a) TPSA is the most important coefficient in the logistic regressor model trained on MPO component values. b) Except for pKa, the coefficients of the logistic regressor tend to be anticorrelated or correlated with TPSA, suggesting they may not add significant BBBP-relevant information beyond TPSA.

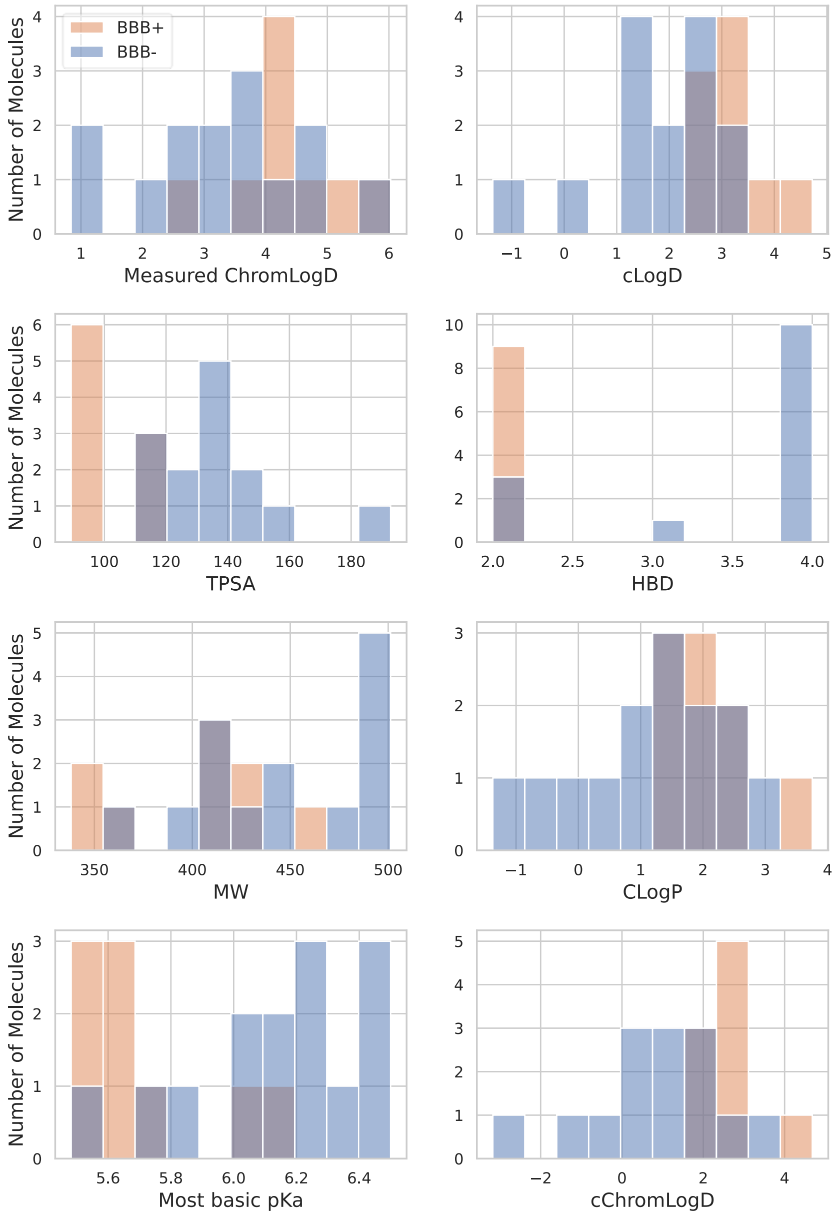


**Figure S7**. Internal Data Distributions of MPO Component Values.

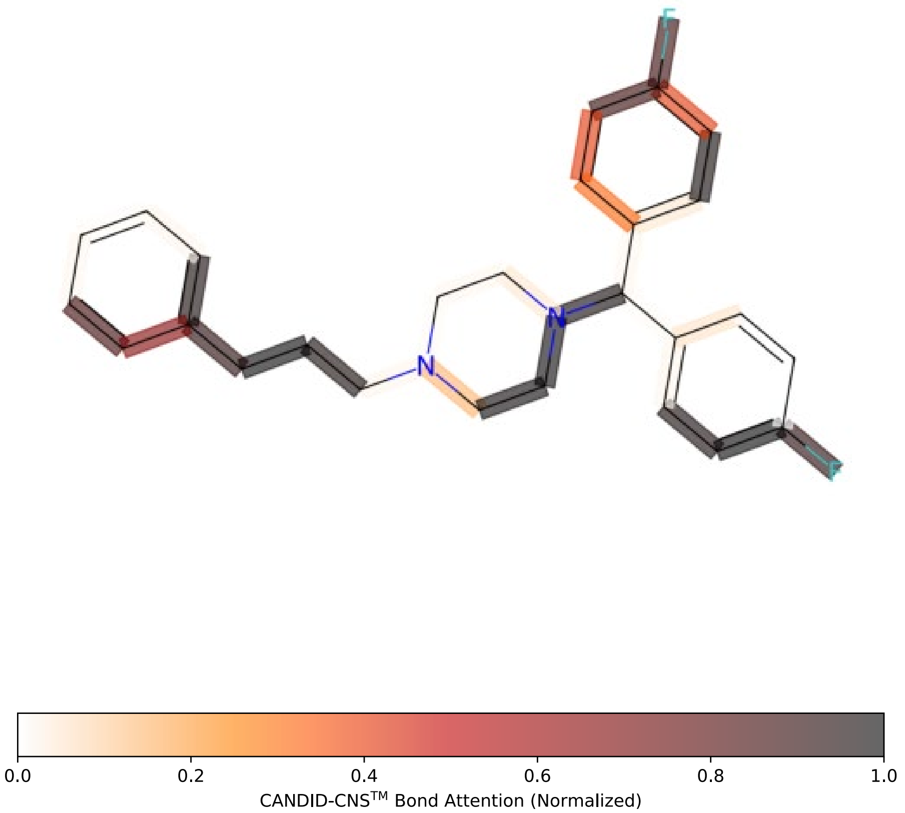


**Figure S8.** Bond Attention Visualization Shows CANDID-CNS™ Learns Insightful Features. Molecule (iv) from figure 5 is visualized here with attention weights. The flexible sp3 carbon bond chain and fluorine atoms are features learned by CANDID-CNS™ that may contribute to this molecule's BBBP.
